# A quantitative and site-specific atlas of the in vivo citrullinome reveals widespread existence of citrullination

**DOI:** 10.1101/2022.12.22.521572

**Authors:** Alexandra S. Rebak, Ivo A. Hendriks, Sara C. Buch-Larsen, Jonas D. Elsborg, Rebecca Kirsch, Nadezhda T. Doncheva, Lars J. Jensen, Maria Christophorou, Michael L. Nielsen

## Abstract

Citrullination is the conversion of peptidyl-arginine into the non-coded amino acid citrulline. Despite its importance in physiology and disease, global identification of citrullinated proteins and precise modification sites has remained challenging. Here, we employed quantitative mass spectrometry-based proteomics to generate a comprehensive atlas of citrullination sites in a physiologically relevant cell type. Collectively, we identified 14.056 citrullination sites within 4.008 proteins and quantified their regulation upon inhibition of the citrullinating enzyme PADI4. Using this rich dataset, we uncover general mechanistic and cell biological principles of citrullination function, while providing site-specific and quantitative information on thousands of PAD4 substrates within cells. Our findings include signature histone marks and numerous modifications on transcriptional regulators and chromatin-related signaling effectors. Additionally, we identify precise citrullination sites on an extensive list of known autoantigens. Collectively, we describe systems attributes of the human citrullinome and provide a resource framework for understanding citrullinaiton at the mechanistic level.

## Introduction

Post-translational modifications (PTMs) are chemical changes that occur on proteins in response to cell stimuli. They are often reversible and act as dynamic molecular switches that modulate protein structure and function, including folding, activity, localization, stability and interaction with other molecules. As such, PTMs provide an enormous amount of complexity to the proteome and enrich the regulatory scope of cells. Deregulation of PTM-catalysing enzymes underlies the development of various diseases and these enzymes are often investigated as key therapeutic targets. As a result, systems-wide characterization of PTMs, as made possible using mass-spectrometry (MS)-based approaches, has become highly relevant for understanding the regulatory mechanisms of protein modifications in human cells and for improving drugs such as protein-based therapeutics^1, 2^.

The advancement of MS-based approaches for studying various PTMs, including lysine acetylation, ubiquitylation, arginine methylation and SUMOylation^3–7^ has significantly improved our understanding of these modifications and provided important advances in molecular and cellular biology^8–10^. However, the cellular extent and biological implications of many protein modifications remain poorly understood.

The conversion of arginine to the non-coded amino acid citrulline, in a process referred to as “deimination” or “citrullination”, is carried out by a small family of enzymes called peptidylarginine deiminases (PADIs, or PADs). The human genome encodes five PADI isoforms, referred to as PADI1, PADI2, PADI3, PADI4 and PADI6, which primarily differ in terms of their tissue and cellular distributions^11^. During the citrullination reaction the guanidinium group of the arginine side-chain is hydrolyzed, removing a positive charge and altering the hydrogen-bonding ability of the modified amino acid. Hence, citrullination can have profound effects on interactions with nucleic acids and other proteins, but may also alter the structure of modified proteins^12–14^.

Citrullination plays functional roles in various biological processes including transcriptional regulation, regulation of pluripotency and reprogramming in the early mouse embryo, immune response to bacterial infections, neuron insulation and skin keratinization and homeostasis ^14–18^. Emerging evidence suggest that citrullination is involved in the etiology of several disorders, with deregulation of PADI activity constituting an abnormal feature in pathologies such as autoimmunity, neurodegeneration, atherosclerosis and cancer^19–21^.

Citrullination in neutrophils is attracting increasing attention as a mediator of rheumatoid arthritis (RA)^22, 23^ and systemic lupus erythematosus (SLE)^24, 25^. In RA, an increase in protein citrullination is observed in the synovial fluid of inflamed joints with >75% of RA patients harboring autoantibodies against citrullinated peptides (CCPs)^26^. Moreover, genetic association studies have identified PADI4 as a susceptibility locus for the development of RA^27–29^. The presence of anti-citrullinated protein antibodies (ACPA) precedes the development of RA and is associated with more severe and faster progressing disease. Therefore, CCPs serve as diagnostic and prognostic clinical markers for RA. Inhibition of PADI activity has shown efficacy in RA, ulcerative colitis, neurodegeneration and cancer disease models^30, 31^ and therefore inhibition of PADIs is emerging as a key therapeutic strategy for these diseases.

Despite great biological and clinical interest in citrullination, our knowledge of *in vivo* citrullination sites and the associated molecular mechanisms remains limited. Hitherto attempts at mapping citrullinated proteins and modification sites have been limited by the analytical strategies used and the ability of mass spectrometers to discern specific, low abundant citrullination events in complex cell extracts. To alleviate some of these challenges, studies have employed cell extracts complemented *in vitro* with active recombinant PADI enzymes^32^. Although providing important knowledge, the citrullination events observed under these experimental conditions may not reflect the *in vivo* biological settings of a living cell, which underscores the need for methodological advances to improve the characterization of citrullination.

We exploited recent advances in proteomics technologies to map citrullination in-depth and at very high accuracy within biological samples. In the current study, we employed powerful high-resolution mass spectrometry, coupled with high-pH fractionation^33^ and computational analysis to define the human citrullinome and to specify the citrullination sites catalyzed by endogenous PADI4. To capture the functional and biological aspects of citrullination, we used the HL60 cell line to model human neutrophil behavior as these cells can be differentiated into neutrophil-like cells (NLCs), resulting in the endogenous expression of PADI enzymes^34^. Increasing the cellular concentration of calcium ions activates the PADI enzymes, which increases catalysis of *in vivo* citrullination events that subsequently can be characterized by quantitative mass spectrometry.

Our proteomic analysis of neutrophil-like HL60 achieves high confidence and allows in-depth characterization of the citrullinome. To pinpoint PADI4-specific citrullination events, we quantified global citrullination changes in response to the PADI4-specific inhibitor GSK484^35^. From this we find that PADI4 targets thousands of non-histone proteins in the nucleus, including a wide range of transcription factors, histone modifying enzymes, chromatin remodeling factors, co-activators and repressors.

Collectively, our analyses identify 14.056 high confident citrullination sites belonging to 4.008 protein coding genes hereby significantly extending the current list of known citrullinated and sites more than 16 fold. Besides pinpointing specific citrullination events on core histone variants, our data describe thousands of non-histone proteins as becoming citrullinated upon PADI activity. Suggesting that citrullination has broad regulatory functions and that the extent of citrullinated autoantigens in neutrophil cells may be much larger than currently anticipated.

Collectively, this work demonstrates that the regulatory scope of PADI4 is much broader than currently anticipated, and comparable in magnitude to important nuclear regulators including the CBP/P300 acetyltransferase ^9^.

## Results

As citrullination occurs with low abundance in cells and constitutes only a small mass change to the modified proteins, it has traditionally been difficult to analyze it using proteomic methods. For confident identification of citrullinated proteins and modification sites on a global scale, we devised a proteomics-based strategy using human promyelocytic leukaemia cells (HL60) in the presence or absence of calcium ionophore. Considering that citrullination sites are often difficult to detect in a large background of non-citrullinated peptides due to their low abundance^36^, we reasoned that pre-fractionation using high-pH reverse-phase fractionation may reduce sample complexity^33^ and improve overall detection efficiency (Figure 1A). In combination with optimized sequencing speed by the MS instrumentation^37^, we surmised that the pre-fractionation strategy would allow direct analysis of citrullination events without the need for a PTM-specific enrichment^38^. To quantify expression changes we employed label-free quantification (LFQ)^39^ which, combined with high-resolution mass spectrometry (MS), would enable identification and quantification of citrullinated peptides with high confidence.

**Figure 1.**
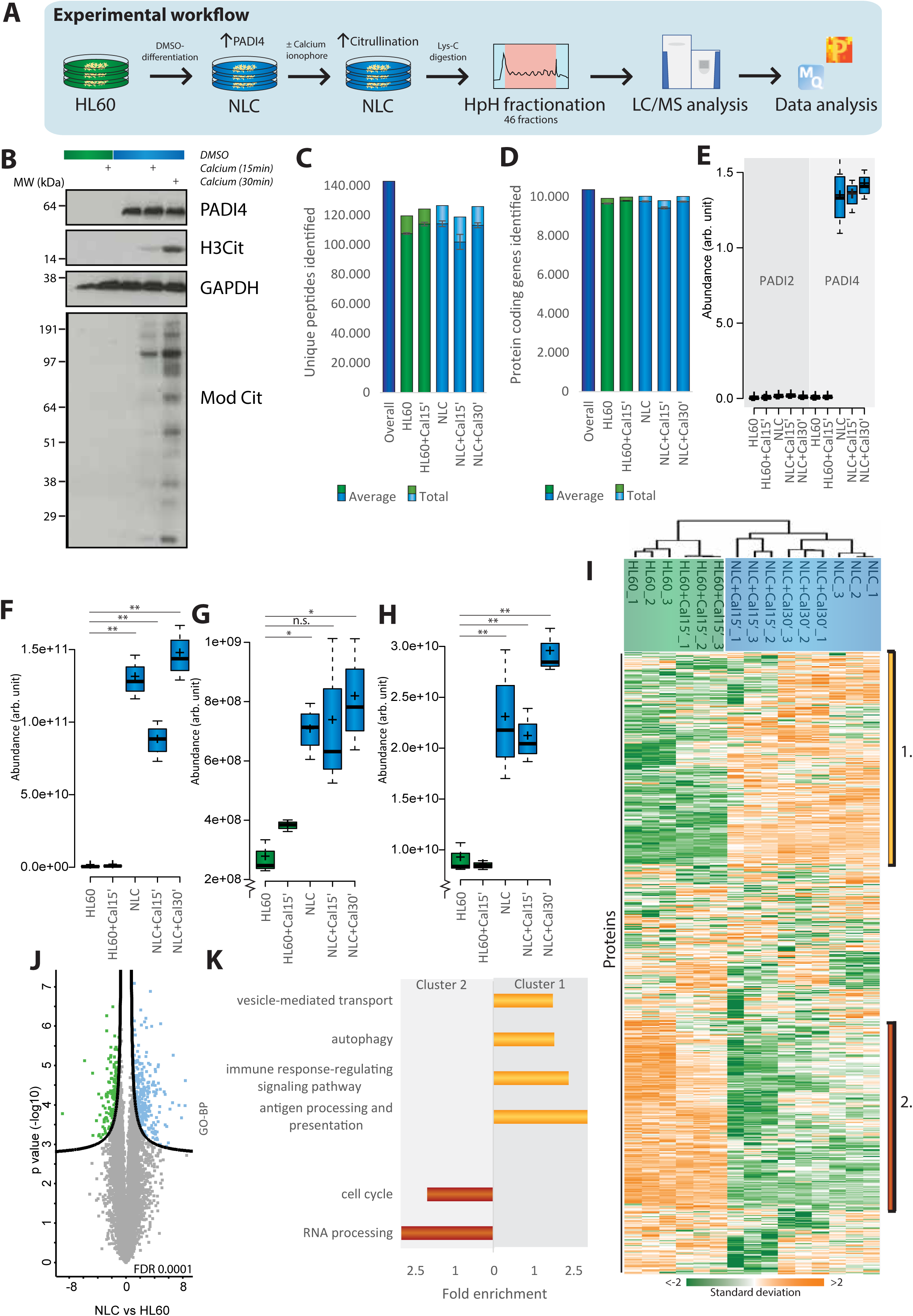
Establishment of regulatory citrullination cell system. **A)** Schematic representation of the workflow. HL60 cells were cultured in standard RPMI medium or differentiated by DMSO to neutrophil-like cells (NLC). Cells were then incubated in Locke’s solution with 4µM calcium ionophore for 0, 15 or 30min. Following Lys-C digestion the peptides were separated by high pH (HpH) fractionation and analyzed using high-resolution LC-MS/MS. Each condition was prepared in triplicates. **B)** Western blot analysis of PADI4, histone 3 R2Cit and modified citrulline shows that PADI4 is expressed upon DMSO differentiation and activated following calcium treatment. **C)** Number of cumulative peptides detected and across the conditions. Error bars represent SD. **D)** Number of cumulative proteins detected and across the conditions. Error bars represent SD. **E)** Abundance of PADI2 and PADI4 across conditions, based on LFQ intensity. Whiskers: 95th and 5th percentile, box limits; 3rd and 1st quartiles, center bar; median, +symbol; average. **F)** Abundance of CD11, **G)** CD16b and **H)** CD55 across conditions. Whiskers: 95th and 5th percentile, box limits; 3rd and 1st quartiles, center bar; median, +symbol; average. Asterisks indicate significant difference between HL60 and the indicated conditions, *P<0.05, **P<0.001, n.s. non-significant. **I)** Hierarchical clustering analysis of z-scored intensity values of all proteins detected across all conditions. (Orange up, green down). **J)** Volcano plot analysis to visualize protein dynamics in response DMSO-differentiation, from HL60 to NLC condition. FDR of 0.0001. **K)** Gene Ontology (GO) term enrichment analysis for biological process of citrullination target proteins of selections displayed in D, as compared to the detected HL60 proteome.

We decided to apply our proteomics strategy to HL60 cells as these can be differentiated into neutrophil-like cells (NLCs) upon provision of retinoic acid or DMSO in culture^34^ leading to increased expression of PADI4. The addition of a calcium ionophore enhances the catalytic activity of PAD4 and since prolonged treatment can recapitulate NET formation *in vitro*^40^ we opted 15 and 30 minutes treatments only.

We confirmed by immunoblot that DMSO-induced differentiation of HL60 into NLCs is accompanied by up-regulation of the PADI4 enzyme (Figure 1B, upper blot). Treating the cells with calcium ionophore leads to activation of the endogenous PADIs and profound global citrullination detectable by immunoblotting including the known citrullination target Histone H3R2^41^ as well as a Pan-peptidyl-citrulline antibody (ModCit) (Figure 1B).

To characterize the proteins that become citrullinated upon PADI activation, we analysed differentiated and undifferentiated HL60 cells (+/- DMSO treatment), in the absence or presence of calcium ionophore. All experiments were performed in triplicate. We used two different calcium ionophore stimulation conditions for differentiated cells, treating for 15 or 30 mins, therefore analysing a total of 5 biological conditions (i) HL60 ÷Calcium; ii) HL60 +15min Calcium; iii) NLCs ÷Calcium; iv) NCLs +15min Calcium; v) NLCs +30min Calcium). To improve identification and localization of arginine residues targeted for citrullination all protein samples were digested with endoproteinase LysC, which cleaves exclusively C-terminal to lysine residues ensuring that arginine residues are positioned internally of the analyzed peptide sequences. For each condition and replicate, we fractionated the samples into 46 fractions using High-pH fractionation as previously described^6, 33^, and analyzed each fraction using 40 min LC-MS gradients on a high-resolution mass spectrometer (Q Exactive HF, Thermo Fischer)^42^. Together we analyzed 138 LC-MS/MS runs for each biological condition, and the combined analysis of the data was performed with the MaxQuant software with a false discovery rate of 1% at the peptide and modification site level.

### Proteome changes upon PADI activation

As we performed HpH-fractionation on whole cell lysate and excluded a PTM-specific enrichment step, our proteomic strategy allows for interrogation of the cellular protein expression changes (i.e. the proteome) simultaneously with the profiling the citrullinome^33^. Overall, the biological replicates of the proteome measurements demonstrated high quantification accuracy and reproducibility with a clear distinction between the two cell populations, HL60 and NLC (Figure S1A) with a Pearson correlation coefficients ranging between 0.97 and 0.99 between replicate analyses (Figure S1B). We identified a total of 143.403 unique peptides at a FDR of 1% across experimental conditions, with individual experiments yielding identification of >100.000 unique peptides (Figure 1C) (Table S1). The identified peptides mapped onto a total of 10.630 unique protein groups and more than 9.400 unique protein groups were quantified across individual experiments (Figure 1D) (Table S2). The HpH-fractionation approach has previously been shown to provide improved peptide sequence coverage of identified proteins ^33^ and we similarly observe that our analysis results in a median sequence coverage of >37% across the identified proteins (Figure S1C).

Consistent with our observations using WB (Figure 1B) and literature reports^34^, our proteome analysis confirms that expression of PADI4 is significantly induced upon the differentiation of HL60 cells into neutrophil-like cells (Figure 1E). Neutrophils are known to express PADI4 and PADI2^43, 44^ and while we found evidence for low expression of PADI2, no expression changes were observed across the investigated cellular conditions (Figure 1E). These observations align with PADI2 being observed across many cell types, whereas PADI4 is more restricted to hematopoietic cells with highest levels in neutrophils ^44^. As our deep proteome analysis were able to profile the protein expression levels for >10.000 proteins across HL60 and NLCs (Figure 1D), we found no evidence for expression of other PADI enzymes beyond PADI4 and PADI2 (Table S2). From this, we conclude that these two enzymes are the likely drivers of citrullination sites induced in the analyzed cells. Besides the increased expression of PADI4, several proteins previously reported as neutrophil markers, including CD11 (Figure 1F), CD16 (Figure 1G) and CD55 (Figure 1H)^45, 46^, were increased in expression upon differentiation into neutrophil-like cells.

Our protein expression data enables a systems-level assessment of DMSO-treatment on the proteome, providing a global protein expression basis for the functional specialization of HL60 cells into neutrophil-like cells. To obtain a functional classification of proteome differences related to differentiation into neutrophil-like cells, we performed unsupervised hierarchical clustering of the >10.000 identified proteins (Figure 1I). The resulting heat map revealed one major cluster of proteins highly expressed in HL60differentiated NLCs (Figure 1I & 1J). Gene ontology analysis revealed that proteins in cluster 1 were enriched (p<0.006) in terms of known neutrophil functions, including immune response-regulating pathways, autophagy, antigen processing and presentation, and vesicle mediated transport (Figure 1K and S1D). The second major cluster (cluster 2) comprised proteins with high expression in undifferentiated HL60 cells, with this cluster enriched in biological processes such as cell cycle and RNA processing. Collectively, we conclude that the differentiation of HL60 cells into NLCs was successful, resulting in the relevant biological conditions for studying citrullination events mediated by the PADI4 enzyme.

### Proteome-wide identification of citrullination in neutrophil cells

Having established NLCs that express PADI4, we evaluated the capability of our methodology to identify citrullination sites. Overall, we observed a high degree of reproducibility across biological repeats of NLCs (Figure S2A), which resulted in ∼70% overlapping citrullinations sites between any two replicate runs of NLCs (Figure S2B). For all investigated cells and treatments a PCA analysis revealed good distinction between the individual cell populations (Figure 2A). Reassuringly, the high precursor mass accuracy of the MS analysis (<2 mDa) ensured that the mass increment caused by citrullination (∼0.9840 Da) can easily be distinguished from the peptide mass delta resulting from the occurring of natural carbon and nitrogen stable isotopes (13C: +1.0034 Da; 15N: +0.9970 Da) ^47^ (Figure 2B & S2C). To pinpoint citrullination sites within each identified peptide we employed high-resolution tandem mass spectrometry (MS/MS) analysis, with each identified spectrum assigned a localization score to depict the confidence in which amino acid in the peptide sequence that carries the citrullination (Figure 2C).

**Figure 2.**
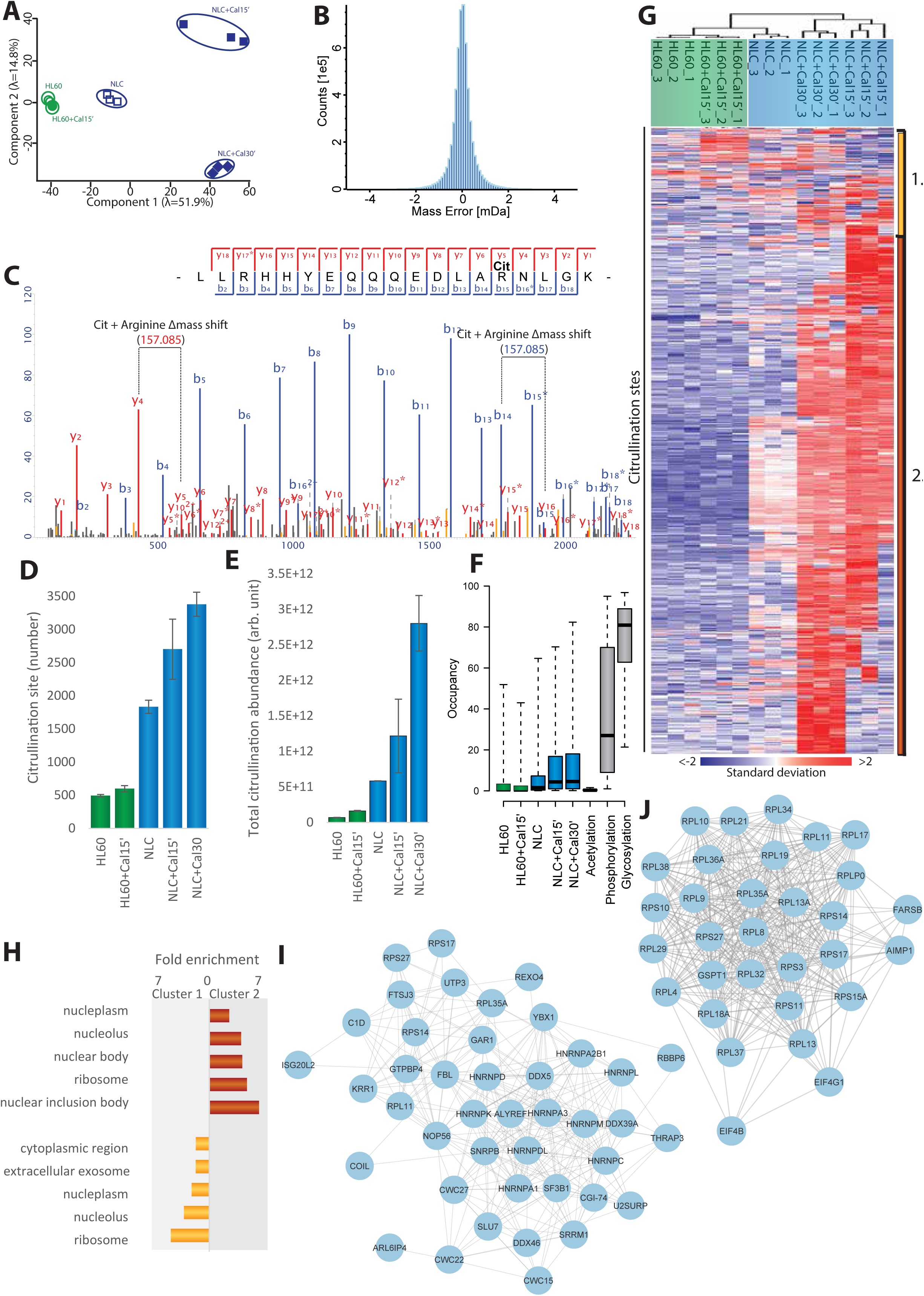
Validation of regulatory citrullination cell system. **A)** Principal component analysis of all conditions at citrullination site level. Eigenvalues are displayed on the axes. **B)** Mass accuracy of all peptides identified. **C)** Representative MS/MS spectrum displaying localized citrulline. **D)** Number of citrullination sites detected in the five conditions by MS/MS and matching. Localization score >0.75. Error bars represent SD. **E)** Total citrullination abundance of sites detected across conditions. Error bars represent SD. **F)** Occupancy of citrullination across the five conditions, acetylation^103^, phosphorylation^104^ and glycosylation^105^. Whiskers: 95th and 5th percentile, box limits; 3rd and 1st quartiles, center bar; median, +symbol; average. **G)** Hierarchical clustering analysis of z-scored intensity values of all proteins detected across all conditions. (Red up, blue down). **H)** Gene Ontology (GO) term enrichment analysis for biological process of citrullination target proteins in HL60 and NLC+Cal30’ condition, in addition to selections displayed in H (cluster 1 and 2), as compared to the human proteome. **I)** STRING-network visualizing functional interactions between proteins annotated to be involved in translation (red) and RNA processing (purple) upregulated in core citrullinome identified across all conditions. STRING clustering score > 0.7 was used and disconnected proteins were omitted from the network.

As citrullination of arginine residues entails the same peptide mass shift as deamidation of glutamine and asparagine residues, we only considered citrullination sites identified from MS/MS spectra entailing high localization probability (>0.9) (Figure S2D), thereby ensuring high confidence in distinguishing between amino acids that become citrullinated or deamidated. Furthering this, the average localization probability of all identified citrullination sites in our filtered dataset was 99.41%.

Collectively, our large-scale analysis resulted in high-confidence identification of 5.238 citrullination sites across the investigated cell conditions (Table S3), including known histone marks and on non-histone targets such as EP300 and NPM1. Within these data, we investigated which proteins were predominantly modified with citrullination (Table 1). The largest number of citrullination sites were observed in NLCs after 30 min stimulation with calcium ionophore (Figure 2D), which correlated with WB observations (Figure 1B). We found >75% of the acquired citrullinated MS/MS spectra (Figure S2F), accounting for ∼95% of the total citrullinated peptide abundance (Figure S2G), to have sustained neutral loss of isocyanic acid, which constitutes a signature for confident identification of citrullination sites^48^. In contrast to the increased citrullination abundance in NLCs (Figure 2E), we observed no significant change in the number of deamidation sites (Figure S2H) across experimental conditions, furthering that of our high-resolution proteomics data facilitates distinguishing between citrullination and deamidation. Notably, we found that the increase in citrullination site abundance between 15 and 30 minutes of calcium ionophore treatment (Figure 2E) is higher than the corresponding increase in the number of citrullination sites for the same conditions (Figure 2D). This indicates that PADI4 activation is specific and tightly regulated, with longer calcium ionophore stimulation leading to enhanced citrullination of the same arginine residues. Overall we are confident that our dataset provides a good coverage of the human HL60 and NLC citrullinome.

Next, we wanted to assess the ‘citrullination modification stoichiometry’ (i.e. the modification percentage of any given arginine residue, also referred to as occupancy), as such information is valuable for understanding the mechanistic implications of PTMs in protein regulation. While stoichiometry by itself does not provide direct evidence for biologically relevant function, it is expected that sites with higher modification stoichiometries are more likely to have functional consequences. Across the citrullinome we observed that the modification stoichiometry increased upon DMSO-induced differentiation and calcium ionophore treatment (Figure 2F). Comparing citrullination stoichiometry to that of other PTMs, found it to be larger than the stoichiometry of acetylation, yet lower than the overall stoichiometry of phosphorylation and N-glycosylation (Figure 2F). Acetylation and phosphorylation are highly reversible PTMs, while glycosylation is thought to be stable and frequently serves structural roles^49–51^. The stoichiometric observations related to citrullination could suggest that it may be reversible, although to date no de-citrullinating enzyme has been described.

### Cellular and Functional Classification of Citrullinated Proteins

To obtain an overview of the sub-cellular compartments and cellular functions that citrulllinated proteins associate with, we first clustered the identified citrullination sites into a heatmap which revealed two main clusters (Figure 2G); one centered upon baseline modification sites (Cluster 1; yellow), and another containing the majority of citrullination sites upon cell stimulation (Cluster 2; red). Next, we performed term enrichment analysis using Gene Ontology (GO) terms to extract the categories overrepresented in the identified citrullination clusters as compared to the human proteome. The GO Cellular Component analysis highlighted that citrullination sites induced upon differentiation into NLCs and subsequent induction of PADI generally localized to the nucleus; and particularly to the nucleolus, nucleoplasm and to proteins associated with the nuclear inclusion bodies (Figure 2H). Modification of the ribosome was also observed which aligns with previous observations^5216^. Although PADI2 has been described to localize to the nucleus upon estrogen and calcium stimulation, PADI4 is generally the only PADI containing a nuclear localization sequence (NLS)^53^, and thus the observed nuclear localization of citrullinated proteins aligns with the localization of PADI4. Citrullination sites that remained unchanged after cellular perturbation (Cluster 1) primarily associated with proteins localized in the cytosol and extracellular exosome, and these sites could be catalyzed by baseline PADI enzyme activity unrelated to the cellular treatments in this study. GO Biological Processes analysis of the two clusters (Figure S2I) indicated an enrichment for protein targeting to ER and glycolysis in Cluster 1, whereas Cluster 2 demonstrated an enrichment for histone modification and proteins involved in regulation of RNA binding.

Unexpectedly, we identified ∼500 citrullination sites in HL60 cells (Figure 2D) not visible by WB analysis (Figure 1B), which suggests low-level activation of PADI enzymes and illustrates the sensitivity of our proteomic approach. The fact that 257 of these citrullination sites were observed across all five conditions (Figure S2J), supports the notion that they may constitute a basal ‘core citrullinome’. The proteins comprising this core citrullinome were significantly enriched for translation and RNA processing (Figure 2I), which suggests that these central cellular functions may be regulated by citrullination under physiological cellular conditions.

### Characterizing the PADI4-specific citrullinome

We next aimed to identify substrates specifically citrullinated by PADI4, by performing quantitative citrullination analysis of cells treated with the PADI4-specific inhibitor GSK484 ^35^ (Figure 3A). To this end, we treated NLCs with GSK484 for 30 min prior to the calcium-induced PADI4 activation, and subsequently compared GSK484-treated cells to mock-treated counterparts with all experiments performed in quadruplicates. To elucidate the concentration-dependent effect of the inhibitor, we performed analysis across three inhibitor concentrations (1uM, 5uM and 20uM) for which strong differences in citrullination were observed on WB (Figure 3B). For the MS analysis we switched to the Orbitrap Exploris™ 480 mass spectrometer for improved MS acquisition^54^. As a result, we observed improved sequencing depth in the samples analyzed (Figure S3A), which overall led to a deeper coverage of the citrullinome (Figure S3B) and improved identification of a total of 14.056 citrullination sites (Table S3) on 4.008 protein coding genes (Table S4).

**Figure 3.**
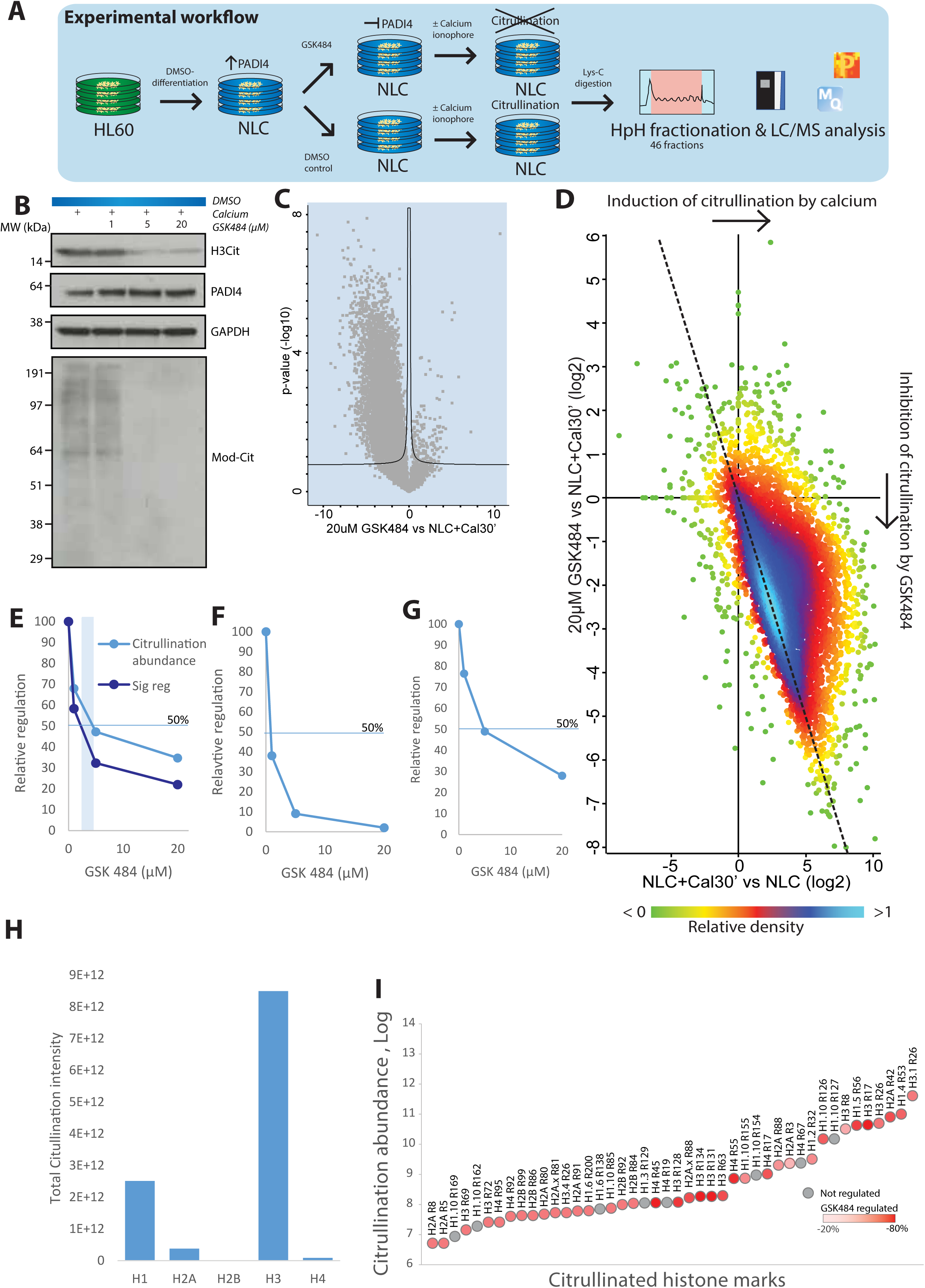
PADI4 inhibition by GSK484. **A)** Schematic representation of the workflow. HL60 cells were cultured in standard RPMI medium and differentiated by DMSO to neutrophil-like cells (NLC). Cells were then incubated in Locke’s solution with varying concentrations of PADI4 inhibitor, GSK484, at 1µM, 5µM, 20µM or a control, for one hour prior to treatment with 4µM calcium ionophore for 30min. Following Lys-C digestion the peptides were separated by high pH fractionation and analyzed using high-resolution LC-MS/MS. Each condition was prepared in quadruplicates. **B)** Western blot analysis of PADI4, histone 3 R2Cit and modified citrulline shows that PADI4 activation by calcium treatment is increasingly inhibited by GSK484. **C)** Volcano plot displaying regulation of sites in response to treatment with 20µM GSK484. **D)** Density plot of sites detected in response to calcium activation (across x-axis) and 20µM GSK484 treatment (y-axis). Dotted line is equivalent to y = - x. **E)** Regulation of total citrullination abundance and significantly regulated sites in response to increasing GSK484 concentration. **F)** Regulation of auto-citrullination site R218 of PADI4 in response to increasing GSK484 concentrations relative to control. **G)** Regulation of citrullination on beta-actin in response to increasing GSK484 concentrations relative to control. **H)** Total citrullination abundance for histone variants. **I)** Diagram showing citrullination abundance of citrullinated histone marks and their level of regulation in response to 20µM GSK484.

Overall, virtually no protein expression level changes were observed when cells were treated with GSK484 (Figure S3C). For the purposes of bioinformatic analysis, we defined significantly regulated citrullination sites based upon significance versus fold-change (Volcano plot) analysis (Figure 3C). Overall, treating cells with GSK484 led to a global reduction in citrullination signal (Figure 3C), and concomitantly a decrease in citrullination site stoichiometry proportionally with increasing concentrations of GSK484 (Figure S3D). To investigate whether citrullination sites induced via calcium activation of PADI4 were inhibited by GSK484, we correlated the increase of citrullination observed upon calcium treatment with the corresponding site-specific changes detected upon GSK484-inhibition (Figure 3D, S3E & S3F). The strong downward correlation between the two datasets confirmed that the majority of citrullination sites induced upon calcium-stimulation of NLCs constitute PADI4 targets. From the Vocano plot analysis (Figure 3C) we found 4.957 citrullination sites as significantly down-regulated upon treatment with GSK484 corresponding to >2.000 citrullination target proteins mediated by PADI4. This corresponded to more than 50% of all identified citrullination target proteins, demonstrating that the regulatory scope of PADI4 is much larger than previously thought. Considering that a hallmark of citrullination-related diseases such as RA are associated with the deregulation of PADI enzymes and subsequent increase in protein citrullination (i.e. autoantigens), we identified citrullination sites on many known ACPA (Table S3) including 44 specifically related to RA based on the Aag Atlas^55^ (Table 2).

As the dynamic turnover rate of citrullination at individual sites remains an almost completely unexplored question, we next exploited the action of GSK484 inhibitor to investigate the kinetics of citrullination on a proteome-wide scale. First, we evaluated the half maximal inhibitory concentration that would be needed to inhibit citrullination events on a global scale. First, we first looked at what concentrations would reduce the total cellular levels of citrullination by 50% (Figure 3E, light blue line). Additionally, as citrullination is regulated at the site level for each substrate, individual proteins harbored modification sites with different citrullination kinetics. To encompass this, we additionally determined when the cumulative signal of all individually inhbited citrullination sites would be reduced in levels by 50% (Figure 3E, dark blue line). Combining both approximations, this yielded a cellular GSK484 IC50 value of 1-3uM, which is roughly 20 times higher than the IC50 reported for GSK484. This difference likely reflects that our analysis was performed in living cells and in the presence of calcium, which is known to reduce the inhibitory effect of PADI4 inhibitors^56^. Moreover, our calculations were based upon the entire proteomics-derived citrullinome whereas standard IC50 values are often based upon low-throughput analysis of a few substrates^35^. Our data furthermore allowed us to look into the site-specific dynamics of citrullination, for example for the known regulatory autocitrullination of PADI4 (R218), which we estimate to have a GSK484 IC50 value of less than 1µM (Figure 3F). Looking at additional autocitrullination of PADI4, we find aberrant citrullination events on the less preferred autocitrullination sites R372 and R374 as previously reported ^57^. However, due to the overall lower abundancy of these modification sites a proper IC50 value could not be established across the investigated cell conditions.indirect

Still, our data shed light on the regulatory effect that PADI4 inhibitors could have on citrullination of proteins commonly citrullinated in RA and multiple schelerosis such as beta-actin (Figure 3G), enolase (Figure S3F) and MBP (Figure S3G)^13, 58, 59^.

Next, we assessed the biological processes associated with PADI4-specific citrullination sites. Performed GO analysis, we identified an enrichment for regulation of RNA export from the nucleus and translational initiation, which matches the core citrullinome (Figure S3I).

### Histone citrullination

PADI4 is a well-established regulator of gene expression through citrullination of histone proteins (Cuthbert, Cell, 2004; Wang, Science, 2004). Additionally, histone hypercitrullination is known to play a critical role in chromatin decondensation in neutrophils^41^. Our data therefore provide a cardinal global survey of the PADI4-mediated histone citrullination events.

First, we confirmed previous observations^60^ that 12-14% of the total cellular citrullination signal in NLCs localizes to histone proteins (Figure S3J), even when the global citrulination signal is decreased upon GSK484 treatment. By contrast, histones only occupy ∼3% of the total protein signal in cells (Figure S3J) demonstrating that core histones are major citrullination targets in human cells. Overall, we observed citrullination sites on all four core histones with highest global citrullination signal observed for Histone H3 (Figure 3H) (Table S4). Generally, the histone citrullination signal increased upon calcium stimulation and was conversely reduced upon PADI4 inhibition, albeit to a differing degree and with different site specificities. The degree of regulation was pronounced for a range of citrullination sites (Figure 3I), with most abundant sites detected on H3, H1 and H2A. For example, citrullination of H3R26, H1R53, H2AR42, and H3R17 sites individually occupying 50%, 13%, 10% and 6% of total histone signal, respectively, with all sites reduced by GSK484 by >80% (Figure 3I) (Table S3). In our analysis we quantified nearly all currently known histone citrullination sites while expanding on the current repertoire, enabling us to obtain a comprehensive map of endogenous PADI4-catalylzed histone citrullination marks (Figure 3I). Although anti-citrullinated H2B antibodies are observed in 90% of RA patients^61^, we find H2B citrullination to only occupy <1% of total histone citrullination levels (Figure 3H), which suggests that even low-abundant histone citrullination marks still entail an antigenic target of the ACPA immune response. The IC50 for H1, H2A and H3 was also estimated (Figure S3K, S3L and S3M).

Despite the broad regulation of histone citrullination, certain marks were reduced to a lesser degree upon GSK484 treatment; for example H3R8, which could indicate these sites may be targeted by additional PADIs or that their turn-over is slower compared to other histone marks such as H3R17 and H3R26. In fact, PADI2 is reported to target same histone marks as PADI4^62^ and while we only observe low cellular expression of PADI2 the enzyme may still be able to catalyze low-level citrullination events. Still, for H3R8 we observed a significant regulation in citrullination upon GSK484 (Table S3), which suggests that PADI4 is the major enzyme responsible for the catalysis of observed histone citrullination sites^63^. Generally, our data supports that PADI4 is able to hypercitrullinate histones^41, 64^ and to a larger degree than previously realized.

Collectively, these results provide a first global map of the PADI4-regulated histone sites in mammalian cells, demonstrating that PADI4 indeed targets all four core histones *in vivo*, but the magnitude of regulation, site-specificity, and inhibition of citrullination are distinct for individual sites and histones. Still, further investigations into the site-specific histone citrullination landscape will enable mechanistic understanding of PADI4-mediated citrullination and transcriptional regulation.

### PADI4 targets transcriptional regulators

Although PADI4 is a known regulator of gene expression and a regulatory function of citrullination of non-histone proteins has been described^65–67^, insights into the site-specific citrullination of transcriptional regulators remains limited. In this dataset we found over 330 transcription factors, chromatin remodelers, histone modifying enzymes and transcriptional co-activators modified with citrullination (Figure 4). For 179 of these we observed increased and, conversely, decreased citrullination upon calcium stimulation and GSK484 treatment, signifying that citrullination of these proteins is mediated by PADI4. Furthering the notion that PADI4 acts at the nexus of a broad range of transcription regulatory pathways.

**Figure 4.**
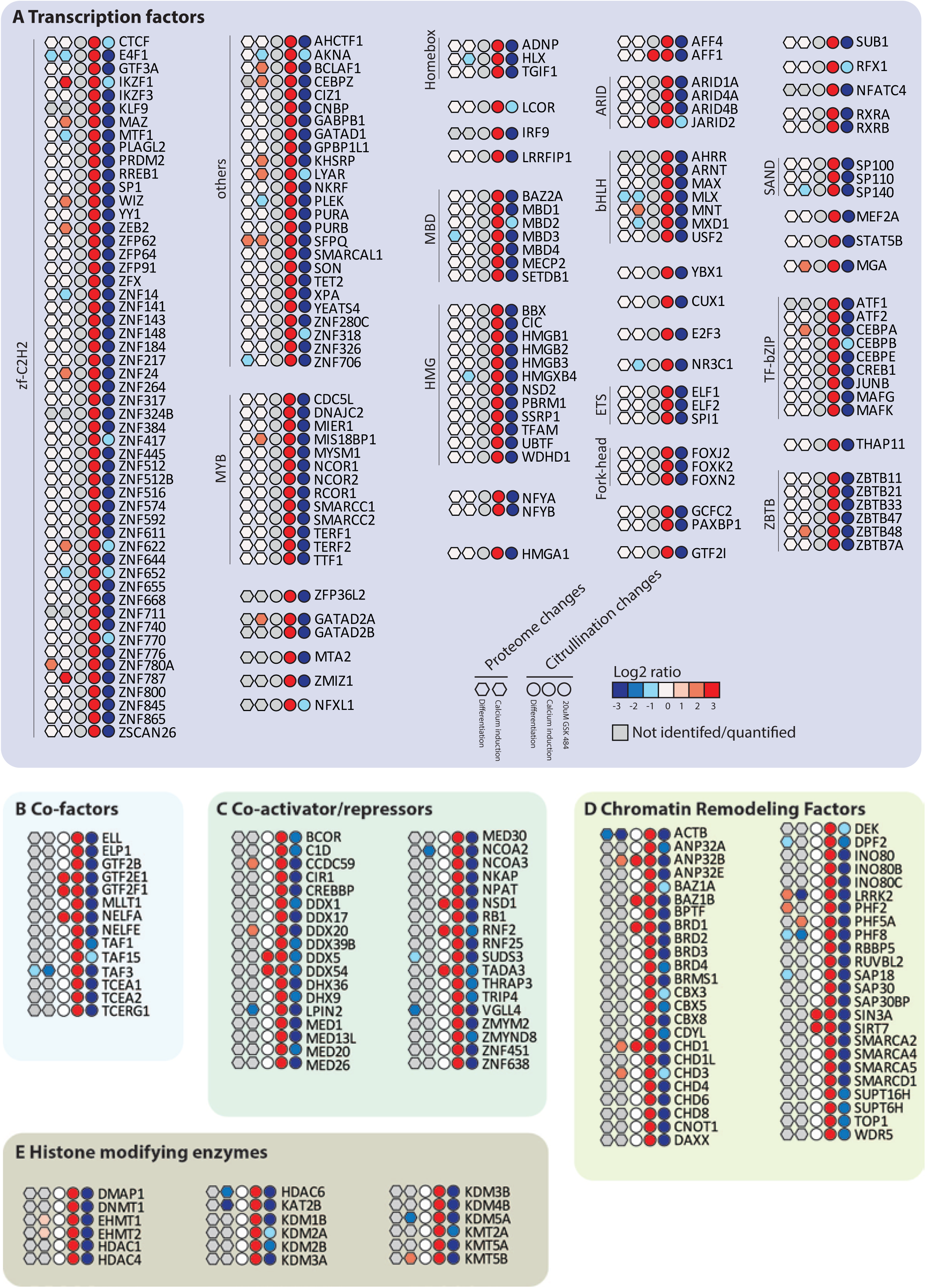
Regulation of transcription factors and other factors by PADI4 activity. Proteome changes in response to DMSO-differentation and calcium induction indicated by color in hexagons for (A) transcriptions factors, (B) co-factors, (C) co-activator/repressors, (D) Chromatin remodeling factors and (E) histone modifying enzymes with significant citrullination regulations indicated by color in circles, either in response to differentiation, calcium induction or 20µM GSK 484.

This wide range of transcriptional regulators within the regulated citrullinome supports previous observations from genome-wide analyses of PADI4 activity, where PADI4 was associated with active genes and as an regulator of gene expression^63, 68, 69^.

Our system-wide approach allows us to consider the downstream targets of the citrullinated transcription factors and investigate which pathways are enriched. While only providing an approximate assessment, we found a high enrichment for pathways related to antimicrobial response and killing of foreign cells, as well as keratinization and cornification (Figure S4). These pathways are known to be linked to citrullination, albeit not at a gene expression level, which provides another level of complexity in understanding the role of citrullination in for example NETosis and myelination. Taken together, these results provides a systems-wide mapping of PADI4-regulated citrullination sites on both histone and non-histone proteins

### Properties of the in vivo citrullinome

Collectively, our large-scale analyses resulted in the high-confidence identification of 14.056 citrullination sites with a localization probability higher than 90% on 3.912 proteins (Figure 5A). Compared to previous *in vivo* studies we expand current knowledge regarding citrullination sites by 16-fold (Figure 5A), and also exceed the number of sites and proteins reported from *in vitro* reaction studies.

**Figure 5.**
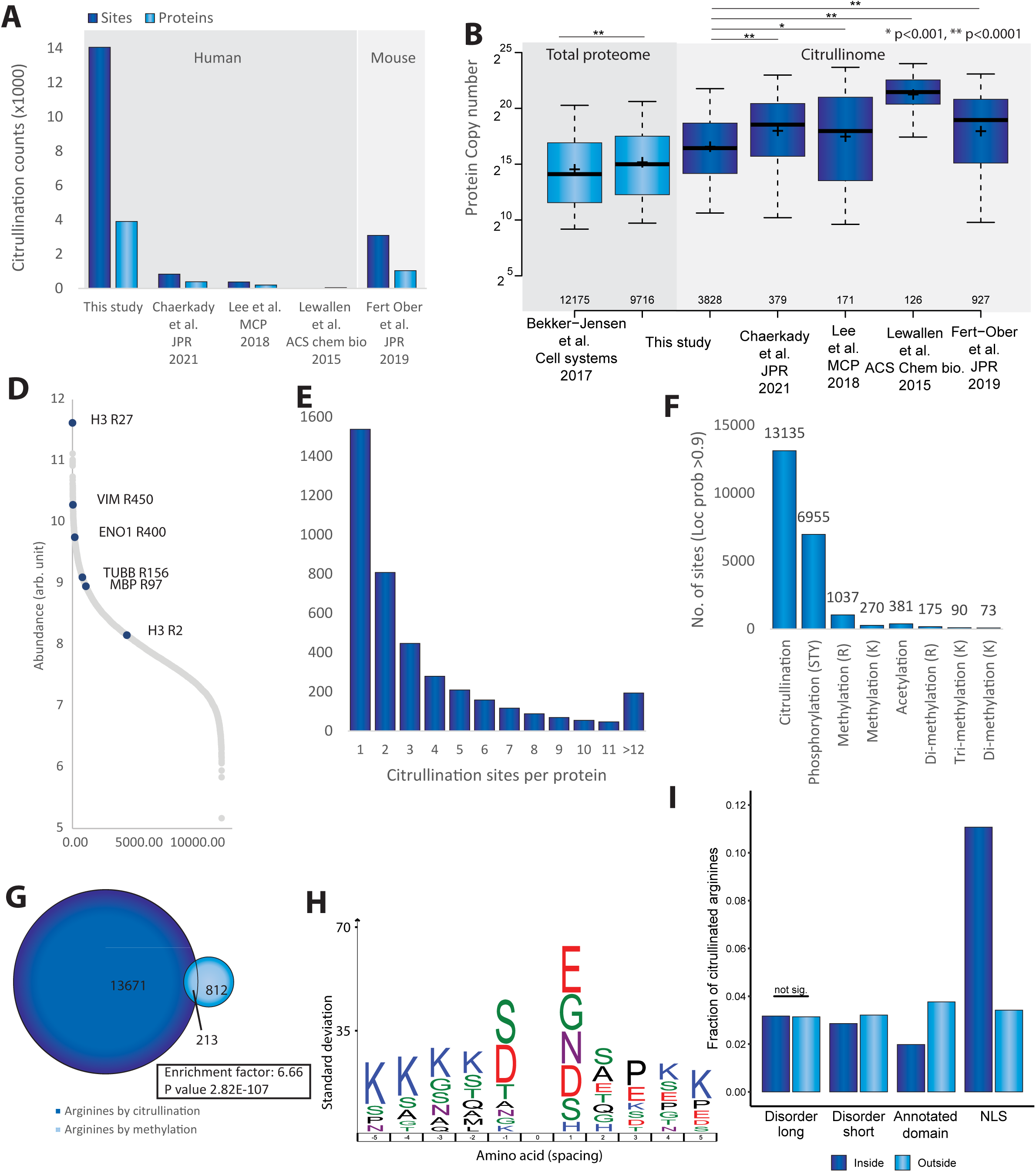
Properties of the citrullinome. A) Number of identified citrullination sites and citrullinated proteins in this study and other citrullination studies. B) Protein copy number of proteomes (light blue) and citrullination screens (dark blue) relative to deep proteome published by Bekker-Jensen et al., 2017. Asterisks indicate significant difference between studies as indicated, *P<0.001, **P<0.0001. C) Distribution of abundance of all sites detected. Most abundant sites on known citrullinated self-antigens indicated across distribution. D) Number of citrullination sites per protein detected. E) Number of common PTMs detected in our dataset F) Pie chart showing the distribution of arginines exclusively targeted by methylation and arginines co-targeted by methylation and citrullination. G) IceLogo representation of enriched amino acid sequence context surrounding identified citrullination sites in DMSO+Cal30’ condition as detected by Exploris, compared to sequence context of all arginines detected in the proteome from the study 1. Amino acid height corresponds to SD change. All displayed amino acids indicate significant changes as determined by two-tailed Student’s t-testing, n=6,245 serine ADP-ribosylation sites, p<0.05. (D) H) Analysis of citrullination sites within different types of protein features show strong enrichment for citrullination sites to localize with nuclear localization signal of proteins.

To assess the depth of sequencing, we compared our study to a comprehensive study of the human proteome. To this end, we plotted the measured iBAQ values of the deep human HeLa proteome and aligned these with the corresponding values for the identified citrullinated proteins and the NLC proteome (Figure 5B). While we achieve comparable depth of sequencing for our NLC proteome analysis, the citrullinome analyses remain less sensitive, which is not surprising considering the overall stoichiometry of citrullination sites (Figure 1F). Still, when assessing the depth of sequencing afforded by our data, the detected citrullination sites span seven orders of magnitude in terms of cellular abundance (Figure 5D), which follows other wide-spread modifications and collectively supports the excellent sensitivity achieved by our proteomics approach.

On average, citrullinated proteins harbored 3.4 citrullination sites, with 40% of proteins detected with only 1 citrullination site (Figure 5E), while 431 proteins harbored >10 citrullination sites, and 26 proteins were citrullinated on >20 arginine residues (Table S4). Hence the distribution of citrullination is relatively similar to other widespread protein modifications such as phosphorylation^4^, arginine methylation^6^ and SUMOylation^7^.

Citrullination and arginine methylation are described to functionally interact on histones in the context of epigenetics and regulation of splicing factors^70, 71^, however, little is known whether the two modifications occupy the same arginine residues within proteins. To assess this, we first analyzed our data for peptides modified with various classes of PTMs, such as phosphorylation, acetylation, as well as various lysine and arginine methylation isoforms (Figure 5F). In total we were able to find 6,955 phosphorylations, 381 acetylations, and 1,645 methylation forms of which the majority (1,037) were identified as arginine mono-methylation sites. Interestingly, 213 citrullination overlapped site-specifically with identified arginine mono-methylation sites, which represented a significantly higher degree of overlap compared to randomly expected (p<2.82E-107). Thus, we hereby show that citrullination and arginine mono-methylation preferentially target the same serine residues on a proteome-wide scale.

Next, we identified a mixed sequence motif for the targeting by PADI4 (Figure 5H), including aspartic acid and serine at position −1 and aspartic acid and glycine at +1 as previously reported ^36^ alongside other amino acids at +1 and −1. The motif is not as strong as detected for other PTMs, however, this is may be due to differences in detection strategies. Where this study is based on a deep citrullinome in a single cell type, Lee et al. 2018 profiled citrullination sites across several human tissues albeit with lower sensitivity.

Considering the functional consequences of the identified citrullination sites, we aimed to test whether citrulination sites localize to specific domains or structural regions of the modified proteins. In this respect, we found that there is an enrichment for citrullination outside short disordered regions and annotated domains (Figure 5I). This matches previous report of citrullination preferentially targeting intrinsically disordered protein regions^12^. Additionally, we identified a strong enrichment for citrullination inside the nuclear localization signal (NLS) of proteins (Figure 5I), along with a predominant targeting of factors related to nuclear processes, including DNA damage, chromatin organization, transcriptional regulation and the cell cycle (Figure 6). This could support an unappreciated role for citrullination in regulating nuclear shuttling and protein localization. Consistent with this notion we found that citrullination of PARP1 (R208), NPM1 (R197) and TDP-43 (R83) are targeted to arginine residues within their respective NLS domains known to regulate the cytoplasmic translocation of the three proteins^72–74^.

**Figure 6.**
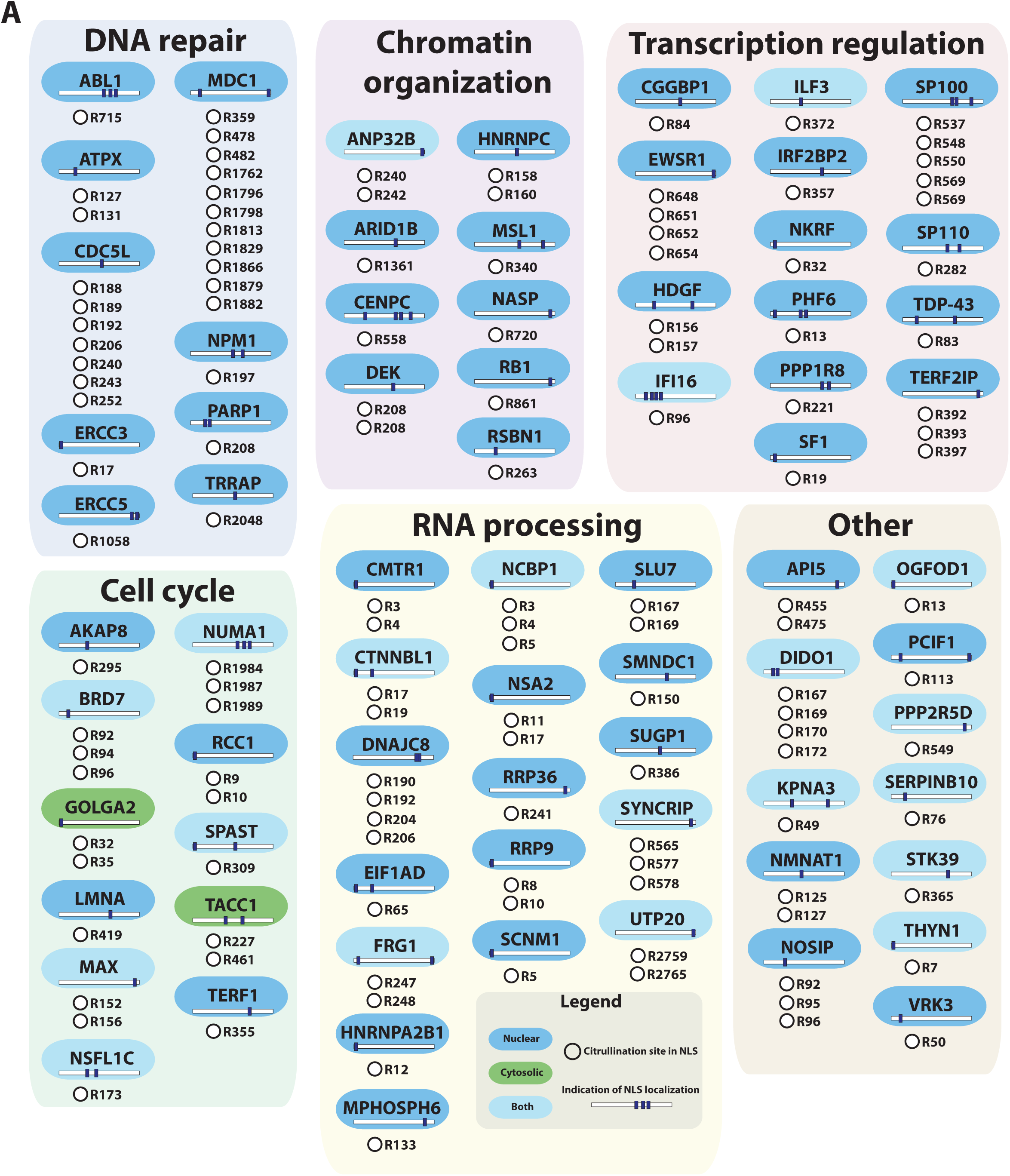
Targeting of nuclear localization signals. A) Representation of proteins significantly targeted by citrullination in NLS and indication of NLS localization in protein sequence. Citrullination site number indicated in addition to potential regulation in response to 20µM GSK48

## Conclusion

Here we present a high-confidence, site-specific atlas including specification of the PADI4-regulated citrullinome. Using high-accuracy mass spectrometry we perform in-depth characterization of over 14.000 citrullination sites from the HL60 model system widely used in the citrullination field. Our data provide insights into the biological and functional implications of citrullination and expand current *in vivo* knowledge regarding this important modification by 16-fold. The ability to pinpoint the residues targeted for citrullination across the proteome hereby provides much needed details regarding its functional consequences, and enables the possibility for developing suitable reagents for biochemical and cell biology studies.

By employing the PADI4 inhibitor GSK488, we demonstrate that the induced sites in our system are specific to PADI4 indicating that the scope of PADI4 targeting is much greater than anticipated. Based upon these observations, our data supports a paradigm for viewing citrullination events mediated by PADI4 as a high-density “citrullination spray” across a multitude of substrates in a manner reminiscent to what has been suggested for other modifications such as SUMOylation^75^ and ADP-ribosylation^76^.

Considering the widespread interest in targeting PADI enzymes to treat a variety of human pathologies^77, 78^, evaluation of the citrullinated proteome under treatment regimes with PADI-specific drugs, as demonstrated in this study, provides insightful information and reveals useful endogenous biomarkers to interrogate pathway function. For example, increasing evidence supports that citrullinated versions of endogenous proteins constitute autoantigens in a variety of autoimmune disorders and that corresponding anti-citrullinated protein antibodies (ACPA) serve as diagnostic and prognostic markers^79, 80^. In support of this, our data details the specific citrullination sites related to 44 autoantigens specific to RA.

Our data also provides insights into the specificity of PADI4 as a peptidylarginine deiminase, and we find that PADI4 citrullinates substrate proteins independently of linear sequence motif and preferentially modifies majority of sites on targeted substrates. We find that citrullination primarily is directed to disordered regions, which supports the role of citrullination in modulating protein binding considering that disordered protein regions hold central roles in protein interaction networks. Similarly, we find that citrullination localize outside annotated domains, except for a strong enrichment of citrullination sites inside nuclear localization signal of proteins. Moreover, we find that PADI4 citrullinates a multitude of transcriptional regulators and enhancer-associated proteins, consistent with its previously reported functions ^69^. GO analysis of the genes regulated by the citrullinated transcription factors shows enrichment for pathways involved in immune responses and terms related to the maintenance of the skin barrier, areas where physiological citrullination is known to play a direct role^41, 81–83^. While these results may only provide indirect clues, they demonstrate the elaborate information entailed in our data and the possibilities for investigating the functional role of citrullination beyond the targeting of individual citrullination sites. While detailed studies are required to understand the regulatory role of PADI4 in gene regulation, our study provides a systems-wide understanding of the relationship between citrullination and gene transcription. Furthering this, we demonstrate that there is a significant overlap in the arginine targeting by citrullination and methylation, supporting previous account on an potential inhibitory interplay between the two PTMs, both known to regulate transcriptional activity^68^

Moreover, histone modifications are widely linked to transcriptional regulation, and great efforts have been made to map the genomic regions targeted by various histone modification marks^84^. For citrullination, however, the understanding of the enzymatic specificity and dynamics remained limited^85^. Within our MS-based quantitative atlas we quantify histone citrullination sites exclusively catalyzed by PADI4, which could help in more accurately defining genomic regions that are occupied by catalytic active PADI4. Especially considering that epigenetic mapping of histone marks typically relates to Histone H3 and H4, and we similarly find that Histone H3 is most abundantly targeted for citrullination by PADI4, indicating that its functional and diagnostic potential may be underappreciated. Collectively, our study provides a first step toward gaining a systems-wide understanding of the complex relationship between PADIs, citrullination, and gene transcription.

Our kinetic analyses upon PADI4 inhibitions provides a first proteome-wide survey of site-specific citrullination events directly mediated via PADI4 via identification of a large set of sites down-regulated upon treatment with GSK484. These regulations are reminiscent of inhibiting other enzymatic regulations within for example phosphorylation and acetylation signaling^10, 86^, and demonstrate that PADI4-mediated citrullination signaling is regulated with a similar degree as other wide-spread moditications.is.

In conclusion, this in-depth analysis represents a powerful resource for the identification and quantification of citrullinated residues induced by PADI4, providing a framework for decoding PADI4 functions in diverse biological processes and a better understanding of PADI4-specific inhibitors, which taken together allows for facile future interrogation of individual site utilization.

## ACKNOWLEDGMENTS

The work carried out in this study was in part supported by the Novo Nordisk Foundation Center for Protein Research, the Novo Nordisk Foundation (grant agreement numbers NNF14CC0001 and NNF13OC0006477), The Danish Council of Independent Research (grant agreement numbers 4002-00051, 4183-00322A and 8020-00220B), and The Danish Cancer Society (grant agreement R146-A9159-16-S2). The proteomics technology applied were part of a project that has received funding from the European Union’s Horizon 2020 research and innovation program under grant agreement EPIC-XS-823839. We would like to thank members of the NNF-CPR Mass Spectrometry Platform for instrument support and technical assistance.

## AUTHOR CONTRIBUTIONS

A.S.R. and M.L.N. designed the experiment. M.C. provided input on project and training on cell handling. A.S.R. prepared the immunoblot. A.S.R. and S.C.B.-L. prepared all MS experiments and A.S.R., S.C.B.-L. and I.A.H. measured all samples on the mass spectrometer and optimised the MS workflow. A.S.R, I.A.H. and J.D.E performed bioinformatics and statistical analyses. N.T.D. and L.J.J. performed transcription factor analysis. R.K and L.J.J. performed structural predictions. M.L.N. supervised the project. A.S.R. and M.L.N. wrote the manuscript with input from all authors.

## DATA AVAILABILITY

The mass spectrometry proteomics data have been deposited to the ProteomeXchange Consortium via the PRIDE^87^ partner repository with the dataset identifier PXD038702

## Methods

### Cell culture

HL60 cells were grown in RPMI media (cat. 21875091, Gibco) supplemented with 10% fetal bovine serum (FBS) and penicillin/streptomycin (100U/mL) (Gibco) at 37°C and 5% CO_2_. A proportion of the cells were differentiated over the course of 4 days to neutrophil-like cells by the addition of 1.25% (v/v) DMSO to the media. The experiments were performed in biological (cell culture) triplicate. Cells were washed in 37°C PBS and transferred to 37°C Locke’s solution (10 mM Hepes pH 7.5, 150 mM NaCl, 5 mM KCl and 2 mM CaCl_2_, 0.1% Glucose) at a concentration of 2×10^6^ cells/mL. Citrullination of proteins was induced via activation of PADI4 by the addition of calcium ionophore Af23187 (cat. C7522, Merck) to a final concentration of 4 µM, for either 15 or 30min, at 37°C. Control samples, of both un-differentiated HL60 cells and differentiated neutrophil-like cells, were harvested from Locke’s solution prior to calcium treatment for further sample preparation.

### PADI4 inhibition by GSK484

Quadruplicate cultures of NLCs were incubated in Locke’s solution, as described above, with a range of GSK484 (cat. SML1658, Merck) concentrations at 37°C for 30 minutes prior to calcium activation, using 4 µM calcium ionophore A23187 (cat. C7522, Merck). The GSK484 concentrations were 1 µM, 5 µM and 20 µM, in addition to a DMSO-control.

### Cell lysis and protein digestion

The cells were pelleted by centrifugation and the Lockes buffer was removed. Cell pellets were then lysed in 10 pellet volumes of Lysis Buffer (6M guanidine-HCl, 50 mM TRIS, pH 8.5). Rapid cell lysis was achieved by alternating vigorous vortexing and vigorous shaking of the samples, 5 seconds per cycle and for 30 seconds in total, after which the lysates were snap frozen in liquid nitrogen. The lysates were stored at −80°C until further sample processing. Lysates were thawed at room temperature (RT) and homogenized using a microtip sonicator, via two pulses of 10 seconds at 30 W. The homogenized lysates were reduced and alkylated, by the addition of Tris(2-carboxyethyl)phosphine (TCEP) and chloroacetamide (CAA), both to a final concentration of 5mM, and incubation for one hour at RT . Proteins were digested using Lysyl Endopeptidase (Lys-C, 1:100 w/w) (cat. 129-02541, Wako Chemicals) for 3 hours at RT. Following a three-fold dilution with 50mM Tris, a second round of Lys-C (1:200 w/w) digestion was performed overnight at RT. Following digestion, samples were acidified via addition of trifluoroacetic acid (TFA) to a final concentration of 0.5% (v/v).

### Purification of peptides

Peptides were purified using reverse-phase C18 cartridges (SepPak Classic, 350mg, Waters). Cartridges were activated with 5 mL acetonitrile (ACN) and equilibrated three times with 5 mL of 0.1% TFA, after which samples were loaded. Sample loading was accelerated using a vacuum manifold maintaining 2/3^rd^ atmospheric pressure. Following loading, cartridges were washed 3 times with 5 mL of 0.1% TFA after which peptides were eluted using 4 mL of 30% ACN in 0.1% TFA. The eluted peptides were frozen overnight at −80°C in 15 mL tubes with small holes punctured into the caps, after which the frozen peptides were lyophilized for 96 hours. Lyophilized peptides were dissolved in 25 mM Ammonium Bicarbonate pH 8.5, and peptide concentration was estimated via absorbance at 280nm, using a NanoDrop instrument.

### Offline high pH reversed-phase HPLC fractionation

For each experimental replicate, 0.6 mg peptide was fractionated into 46 fractions using an XBridge BEH130 C18 3.5µm 4.6mm x 250 mm column (Waters) on an Ultimate 3000 HPLC system (Dionex), operating at a flow rate of 1 mL/min. The flow was composed of three buffers; Buffer A (Milli-Q water), buffer B (100% ACN, and buffer C (25 mM ammonium hydroxide). Prior to loading, samples were basified to pH >10 by the addition of ammonium hydroxide. Peptides were loaded onto the column at 1 mL/min for 4 min, after which they were separated on a linear gradient ranging from 5%B to 25%B over 62 min. followed by an increase to 60 %B over 5 min and 70%B over 3min. Fractions were collected into a 96-deep well plate every 60 to 90 seconds. Following the primary gradient, fraction collection was stopped and the column flow was kept at 70%B for an additional 5 min before it was reduced to 5%B. Buffer C was constant throughout the gradient at 10%. Collected fractions were transferred to Eppendorf Protein LoBind tubes with small holes punctured in the caps, and frozen at −80°C overnight. The frozen samples were lyophilized for 96 hours and afterwards dissolved in 1% formic acid (FA).

### Mass spectrometry analysis

Samples were measured using a Q-Exactive HF-X mass spectrometer or an Exploris 480 mass spectrometer (Thermo Fisher Scientific). Peptides were separated by online reversed-phase liquid chromatography using an EASY-nLC 1200 system (Thermo), using a 15 cm long analytical column, with an internal diameter of 75 µm, packed in-house using ReproSil-Pur 120 C18-AQ 1.9 µm beads (Dr. Maisch). The analytical column was heated to 40°C using a column oven, and peptides were eluted from the column using a gradient of buffer A (0.1% FA) and buffer B (80% ACN in 0.1% FA). The gradient ranged from 4% to 38% buffer B over 30 minutes, followed by an increase to 90% buffer B over 4 minutes to ensure elution of all peptides, followed by a washing block of 6 minutes.

For the pilot experiment, performed on the Q-Exactive HF-X instrument, electrospray ionization was achieved using a Nanospray Flex Ion Source (Thermo). Spray voltage was set to 2 kV, capillary temperature to 275°C and RF level to 40%. Full scans were performed at a resolution of 60,000, with a scan range of 300 to 1,750 m/z, maximum injection time of 60 ms, and an automatic gain control (AGC) target of 3,000,000 charges. Precursors were isolated at a width of 1.3m/z, with an AGC target of 200,000 charges. Repeated sequencing of selected precursors was excluded by dynamic exclusion of 60s. Precursor fragmentation was achieved using higher energy collision dissociation (HCD). MS/MS were measured using the Orbitrap with a maximum injection time of 90ms and a resolution of 45,000. Top9 data-dependent MS/MS method was used to acquire MS data.

For the optimized GSK experiment, performed on the Exploris 480 instrument electrospray ionization was achieved using a NG Ion Source (Thermo). Spray voltage was set to 2 kV, capillary temperature to 275°C and RF level to 40%. Full scans were performed at a resolution of 120,000, with a scan range of 300 to 1,750 m/z, maximum injection time set to “Auto”, and normalized AGC target was set to “200” (2,000,000 charges). Precursors were isolated at a width of 1.3m/z, with a normalized AGC target of “200” (200,000 charges). Repeated sequencing of selected precursors was excluded by dynamic exclusion of 60s. Precursor fragmentation was achieved using HCD. MS/MS were measured using the Orbitrap with a maximum injection time set to “Auto” and a resolution of 30,000

### Western blotting

Cell pellets were lysed in SDS Lysis Buffer (2% SDS, 50 mM Tris-HCl pH 8.5, 150 mM NaCl), and homogenized by heating to 99°C and shaking at 1,400 RPM for 30 minutes. Protein concentrations across lysates were equalized using the Pierce BCA Protein Assay Kit (cat. 23225, Pierce) according to the manufacturer’s instructions. Immublot was performed using standard approaches on an Invitrogen chamber and blot module. Proteins were transferred to PVDF membrane (Immobilon) for one hour and 30 min at 0.4A. Membranes were blocked using 5% milk (Fluka® Analytical) in PBS supplemented with Tween-20 (0.1%; PBST) or 5% bovine serum albumin (BSA) in PBST, depending on recommendations by antibody manufacturer. The following anti-bodies were used: rabbit polyclonal PADI4 (cat. P4749, Sigma Aldrich) and rabbit monoclonal H3 (citrulline Arg2) antibody (cat. Ab176843, Abcam). The Anti-Citrulline (Modified) Detection Kit (cat. 17-347B, Merck) was used to measure global citrullination ^88^, according to the manufacturer’s instructions.

Global citrullination was detected with the antimodified citrulline (AMC) Detection Kit (cat. 17-347B, Merck) according to the manufacturer’s instructions.

### Data analysis

The raw mass spectrometry data files were analyzed using MaxQuant software (version 1.5.3.30), a freely available software routinely used in the field ^89, 90^. Default MaxQuant setting were used with exceptions outlined below. A HUMAN.fasta database was extracted from UniProt on May 5, 2020 to serve as a theoretical spectral library. The HUMAN.fasta database contained 96,821 protein entries. Enzyme cleavage specificity was set to Lys-C. Protein N-terminal acetylation, oxidation (M), phosphorylation (S, T, and Y), deamidation (N, Q, and R), methylation (K and R), di-methylation (K and R), tri-methylation (K), and acetylation (K), were all included as variable modifications. Citrullination, i.e. deamidation of R, was further defined with expected neutral loss of cyanic acid (HNOC, 43.01 Da), and with the immonium ion as diagnostic peak (H_11_C_5_N_3_O, 129.09 Da). A maximum of 3 variable modifications per peptide and a maximum of 2 missed cleavages were allowed. Matching between runs was enabled, with a match window of 0.7 minutes and an alignment time window of 20min. Default settings for filtering by posterior error probability were used to achieve a false discovery rate of <1% at the peptide-spectrum match, protein assignment, and site-specific levels. Label free quantification and iBAQ were enabled.

### Data filtering

In addition to automatic filtering and FDR control as applied by MaxQuant, the data was manually filtered to ensure proper identification and localization of citrullination. Citrullination-site identification was only allowed if localization probability was >0.90. For quantification of citrullination, further PSMs were accepted with localization of >0.75. MaxQuant intensity values (a quantitative metric corresponding to peak area-under-the-curve at the MS1 level) were manually mapped to citrullination sites based on >0.75 localized PSMs only.

### Statistical analysis

Statistical handling of the data and hierarchal clustering was primarily performed using the freely available Perseus software (version 1.6.14.0) ^91^. Significantly enriched Gene Ontology terms were determined using the Functional Annotation Tool of the DAVID Bioinformatics database ^92, 93^. Venn diagrams were generated using the online DeepVenn program ^94^. Boxplots were generated using the BoxPlotR web tool ^95^. Kinase-substrate relationships were predicted using the online NetworKIN tool ^96, 97^. The sequence motif was generated using the iceLogo software (version 1.2)^98^, with background sequences extracted from non-citrullinated arginine residues in all citrullinated proteins.

### Analysis of transcription factor target genes and gene-set enrichment

For each citrullinated and non-citrullinated transcription factor (TF), all target genes were retrieved from TFEA.ChIP^99^ using the ReMap 2020 & GeneHancer Double Elite dataset. Data was available for 115 citrullinated TFs and 233 non-citrullinated TFs. For each of the 16,544 target genes with an Ensembl annotation, the fraction of citrullinated and non-citrullinated TFs, which regulate it, was determined (Table S6). The final score was calculated as the logarithm of the ratio of citrullinated versus non-citrullinated fractions. This value is negative for a given target if there are more non-citrullinated TFs regulating the target than citrullinated ones, and vice versa. All targets were ordered by this value and the whole ranked list was used as input for the STRING v11 gene-set enrichment analysis^100^. The resulting enriched annotations with a false discovery rate below 0.05 are provided in Table S7. A given annotation describes the citrullinated TFs if it is enriched at the bottom of the input, and the non-citrullinated TFs if it is enriched at the top of the input.

### Enrichment analysis of regions targeted for citrullination

For the analysis of citrullination sites in disordered regions, disorder was predicted using IUPred2A^101^ for all identified sequences. The set of long disordered regions was obtained using IUPred2A’s long disorder option and a minimum region length of 31 consecutive residues with a prediction score >= 0.5. Regions predicted using the short disorder option were retained if they contained 2 to 30 consecutive residues with a score >= 0.5. Predictions from all identified sequences were included in the analysis of disordered regions.

Annotations of domains and nuclear localization signals (NLS) were obtained in gff format from UniProtKB ^102^ and their sequences derived from the sequences identified in the MS run. All identified sequences with any kind of feature annotation were included in the domain and NLS analyses.

For each feature category (long and short disorder, domain, NLS), Fisher’s Exact Testing was performed using the counts of unmodified arginine residues and citrullinated arginine residues inside and outside the features. Fold enrichment of citrullines inside features compared to citrullines outside features was calculated as

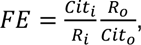

where Cit_i_ and Cit_o_ refer to citrullines inside and outside features, respectively, and R is the total number of unmodified and modified arginine residues.

## Supplemental data legends

**Supplemental Table 1**

A list of all peptides identified in the pilot study (study 1) on the HF-Exactive system and the inhibitor study (study 2) performed on the Exploris system.

**Supplemental Table 2**

A list of all protein coding genes identified

**Supplemental Table 3**

A list of all unique identified citrullination sites in the pilot study (study 1) on the HF-Exactive system and the inhibitor study (study 2) performed on the Exploris system.

**Supplemental Table 4**

A list of all citrullination targeted protein coding genes identified

**Supplemental Table 5**

Statistical information relating to term enrichment analysis, as found in figure 1, 2, S2 found in separate tabs in the Excel file.

**Supplemental Table 6**

Target genes ranked by their relative regulation by citrullinated or not-citrullinated TFs (input file for STRING gene-set enrichment uses column log_ratio_fraction and ensembl_gene_id).

**Supplemental Table 7**

Results from STRING gene-set enrichment for all categories of functional annotations.

---

**Supplemental figure 1.**
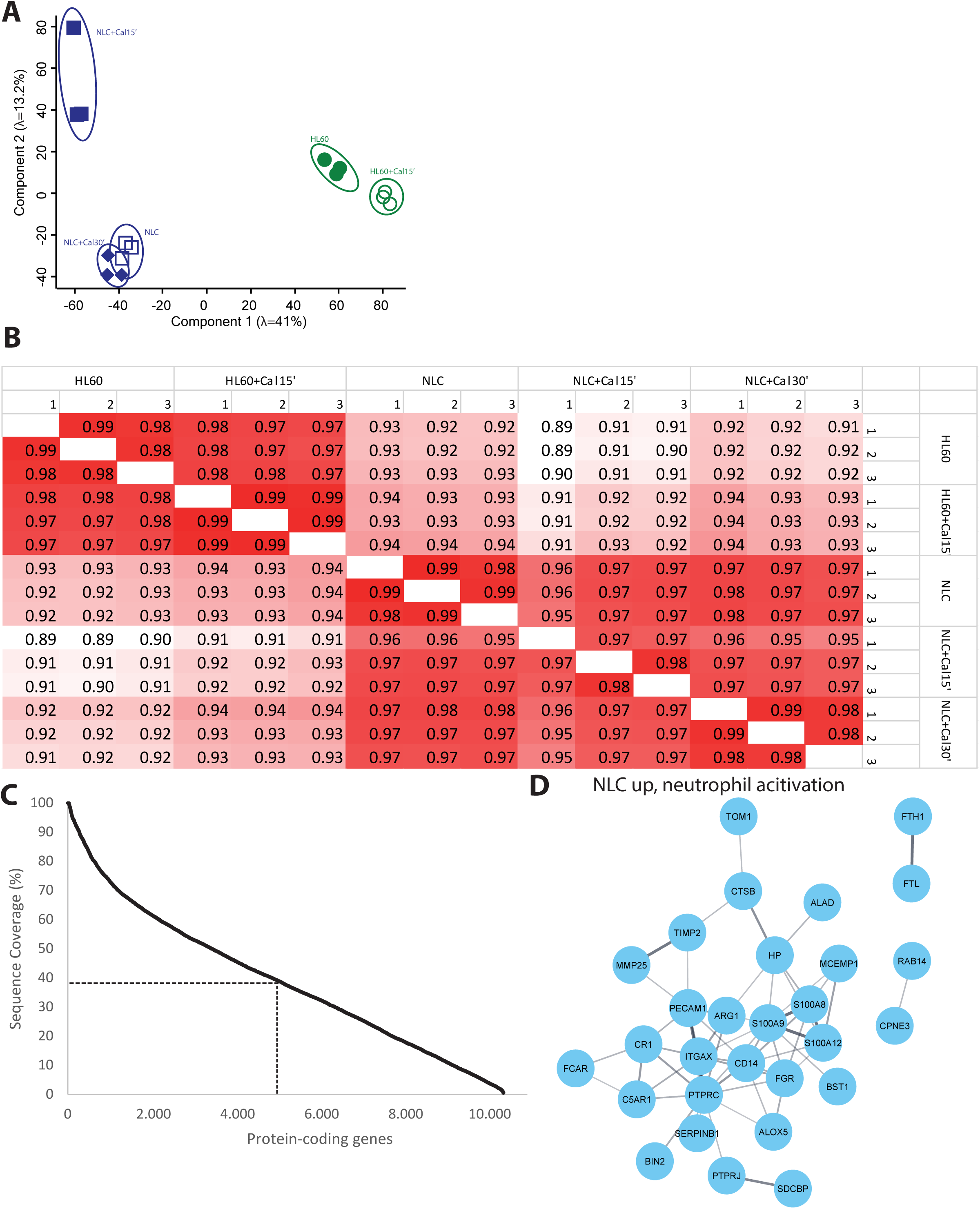
**A)** Principal component analysis of all conditions at protein level. Eigenvalues are displayed on the axes. **B)** Pearson correlation of proteins detected in all replicates of experimental conditions. Clear distinction detection in proteome following DMSO-differentiation. **C)** Sequence coverage of proteins detected. Dotted line indicates median. **D)** STRING network visualizing functional interactions between proteins annotated to be involved in antigen processing and presentation detected as upregulated in NLCs (Cluster 1, Figure 1I & Figure 1K). Default STRING confidence score > 0.4 was used.

**Supplemental figure 2.**
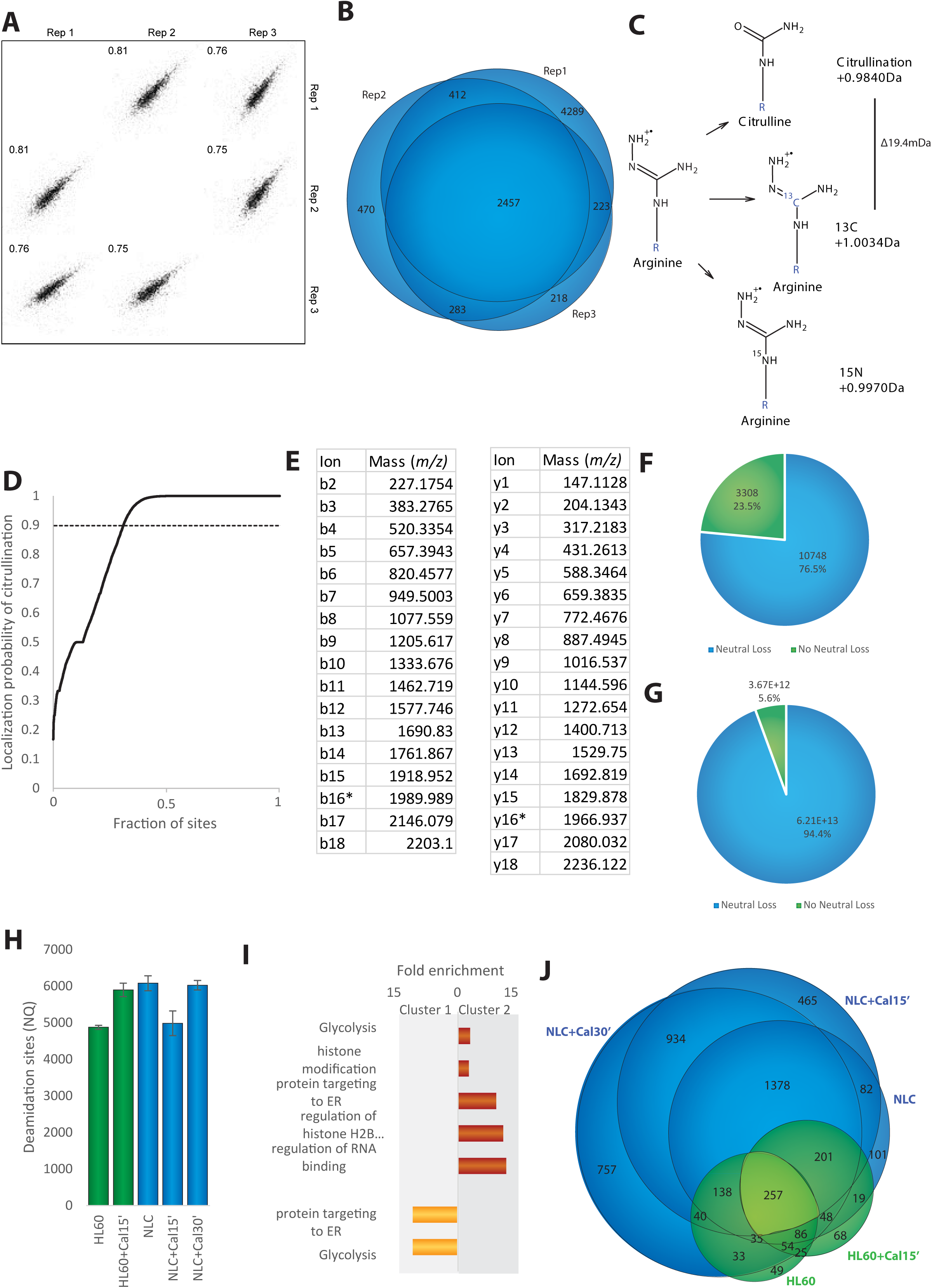
**A)** Pearson correlation of citrullination sites detected in all three replicates of NLC+Cal30’ condition. **B)** Venn diagram showing site detection overlap between all three replicates of NLC+Cal30’ condition. **C)** Diagram displaying small mass shifts between citrullination, carbon-13 (13C) and nitrogen- 15 substitution on residue. **D)** Citrullination localization probability plotted against the ranked fraction of all peptide-spectrum-matches (PSMs). Despite all probabilities being displayed, only those above 0.9 were used for assignment of unique citrullinated peptides and sites. **E)** Mass of b and y ions shown in MS/MS spectrum in figure 2C. **F)** Number of deamidation sites of asparagine (N) and glutamine (Q) detected across conditions. Error bars represent SD. **G)** Pie chart showing distribution of sites detected with neutral loss of cyanic acid or no neutral loss. **H)** Pie chart showing citrullination abundance associated with sites for which a neutral loss was either detected or not detected. **I)** Gene Ontology (GO) term enrichment analysis for cellular components of citrullination target proteins in selections displayed in 1G (cluster 1 and 2), as compared to the human proteome. **J)** Scaled venn diagram displaying overall of sites detected in each experimental condition.

**Supplemental figure 3.**
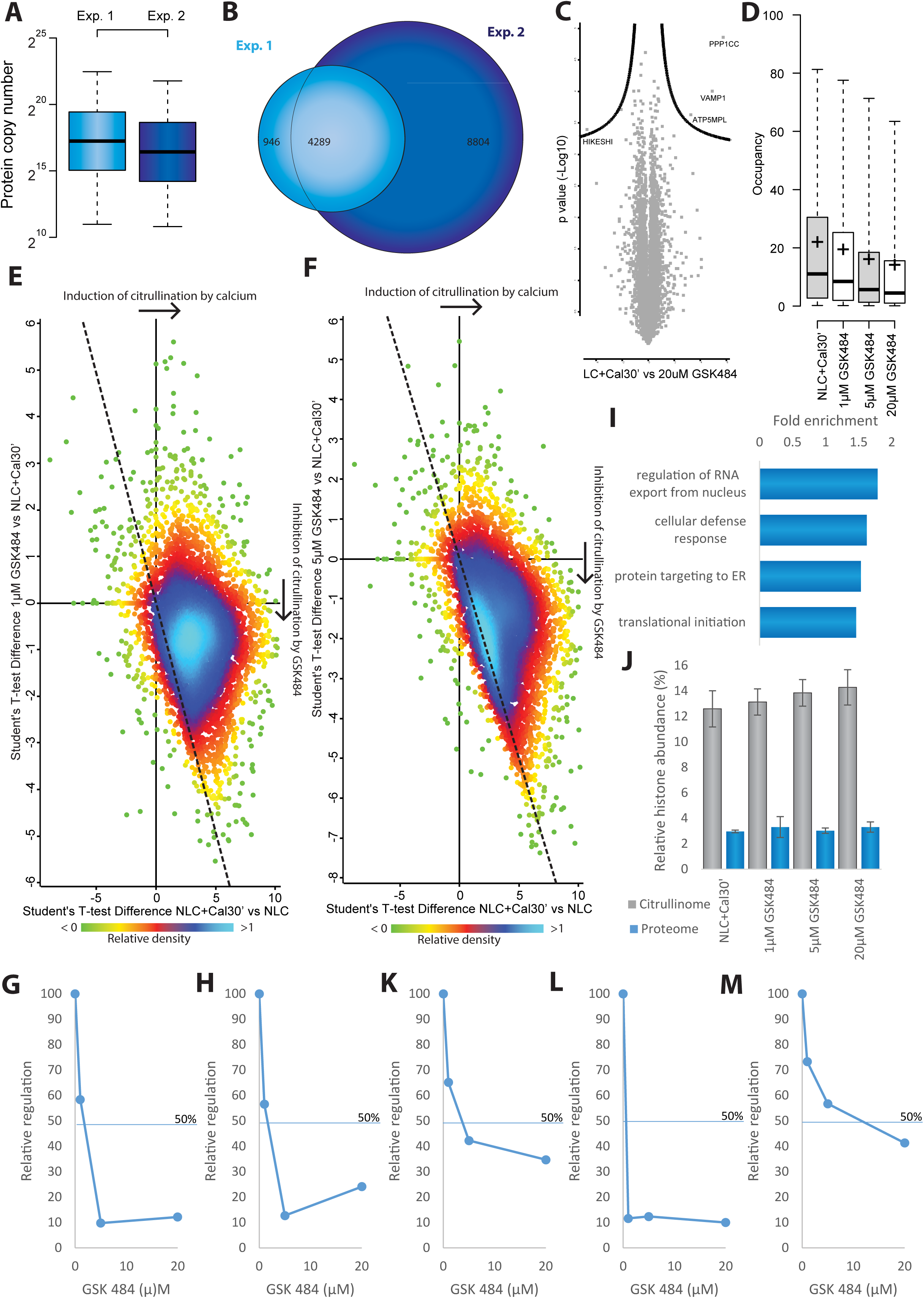
**A)** Protein copy number of proteins detected in pilot study (figure 1 and 2) and GSK experiment with optimized deeper sequencing. **B)** Venn diagram displaying distribution of sites detected in pilot study (figure 1 and 2) performed on HF-Exactive and optimized GSK experiment performed on Exploris systems. **C)** Volcano plot displaying little significant protein regulation in response to treatment with 20µM GSK484. **D)** Occupancy of citrullination across the four conditions. Whiskers: 95th and 5th percentile, box limits; 3rd and 1st quartiles, center bar; median, +symbol; average. **E)** Density plot of sites detected in response to calcium activation (across x-axis) and 1µM GSK484 treatment (y-axis). Dotted line is equivalent to y = - x. **F)** Density plot of sites detected in response to calcium activation (across x-axis) and 5µM GSK484 treatment (y-axis). Dotted line is equivalent to y = - x. **G)** Regulation of citrullination on enolase in response to increasing GSK484 concentrations relative to control. **H)** Regulation of citrullination on MBP in response to increasing GSK484 concentrations relative to control. **I)** Gene Ontology (GO) term enrichment analysis for biological processes of citrullination targeted proteins in the 20µM GSK484 condition, as compared control condition without inhibitor. **J)** Relative histone abundance in citrullinome (grey) and proteome (blue). **K)** Regulation of citrullination on histone 1 variant in response to increasing GSK484 concentrations relative to control. **L)** Regulation of citrullination on histone 2A in response to increasing GSK484 concentrations relative to control. **M)** Regulation of citrullination on Histone 3 in response to increasing GSK484 concentrations relative to control.

**Supplemental figure 4.**
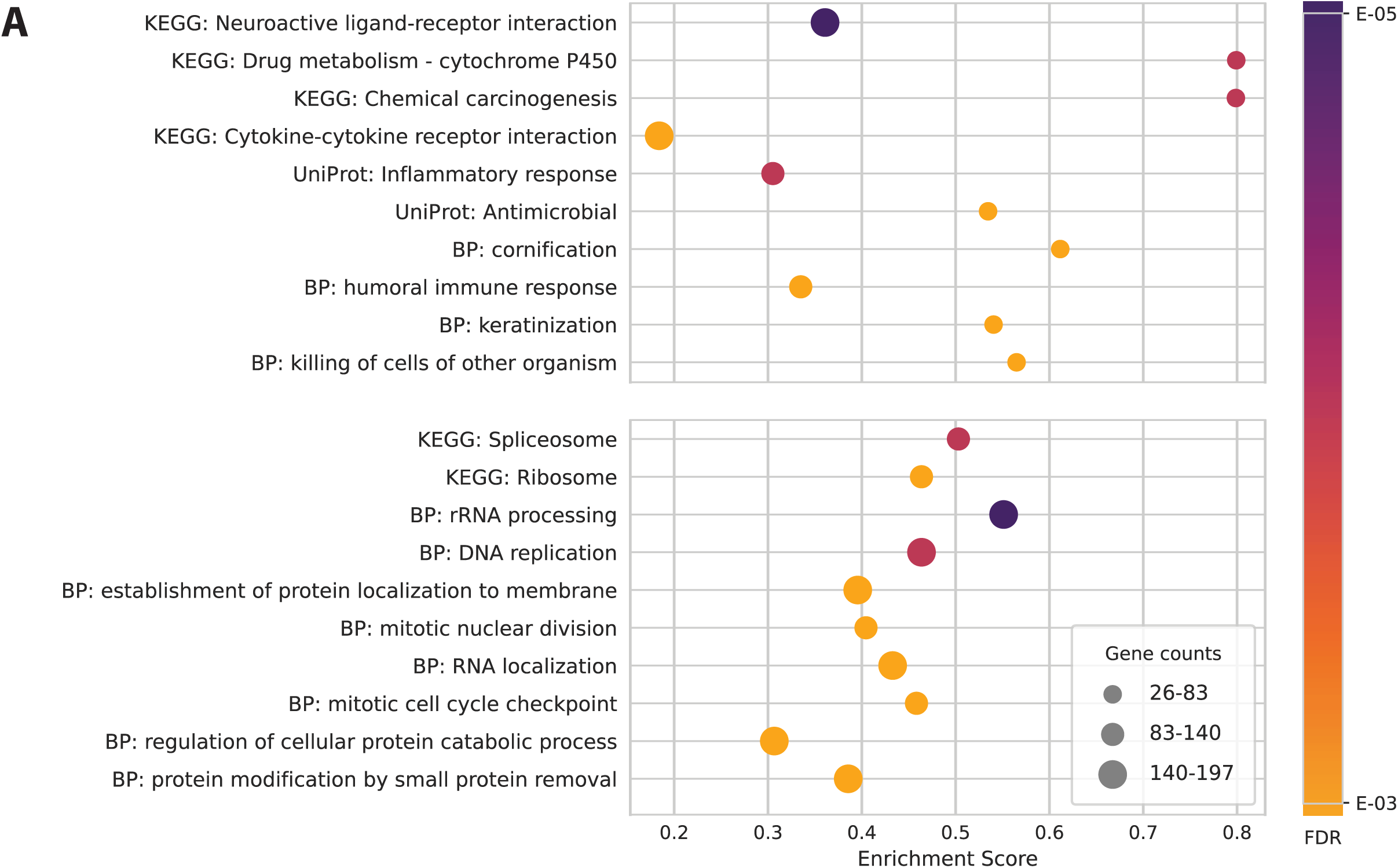
Annotations enriched for the targets of citrullinated (top) and not citrullinated TFs (bottom) from the categories KEGG pathways, Gene Ontology Biological Process (BP), and UniProt keywords as determined by STRING gene-set enrichment analysis (see Table S6 & S7) for a list of all enriched annotations). Gene count refers to the number of genes annotated with the given annotation term, the false discovery rate (FDR) indicates the significance for each term, and enrichment score is the ratio between the term mean and the maximum deviation from the mean in the user input, multiplied by a factor of 10.

## Bibliography

1. Choudhary, C. & Mann, M. Decoding signalling networks by mass spectrometry-based proteomics. Nat. Rev. Mol. Cell Biol. 11, 427–439 (2010).

2. Budayeva, H. G. & Kirkpatrick, D. S. Monitoring protein communities and their responses to therapeutics. Nat. Rev. Drug Discov. 19, 414–426 (2020).

3. Choudhary, C. et al. Lysine acetylation targets protein complexes and co-regulates major cellular functions. Science (80-.). 325, 834–840 (2009).

4. Huttlin, E. L. et al. A tissue-specific atlas of mouse protein phosphorylation and expression. Cell 143, 1174–1189 (2010).

5. Kim, W. et al. Systematic and quantitative assessment of the ubiquitin-modified proteome. Mol. Cell 44, 325–340 (2011).

6. Larsen, S. C. et al. Proteome-wide analysis of arginine monomethylation reveals widespread occurrence in human cells. Sci. Signal. 9, (2016).

7. Hendriks, I. A. et al. Site-specific mapping of the human SUMO proteome reveals co-modification with phosphorylation. Nat. Struct. Mol. Biol. 24, 325–336 (2017).

8. Ordureau, A., Münch, C. & Harper, J. W. Quantifying Ubiquitin Signaling. Mol. Cell 58, 660– 676 (2015).

9. Weinert, B. T. et al. Time-Resolved Analysis Reveals Rapid Dynamics and Broad Scope of the CBP/p300 Acetylome. Cell 174, 231–244 (2018).

10. Ochoa, D. et al. The functional landscape of the human phosphoproteome. Nat. Biotechnol. 38, 365–373 (2020).

11. Vossenaar, E. R., Zendman, A. J. W., van Venrooij, W. J. & Pruijn, G. J. M. PAD, a growing family of citrullinating enzymes: genes, features and involvement in disease. BioEssays 25, 1106–1118 (2003).

12. Tarcsa, E. et al. Protein unfolding by peptidylarginine deiminase. Substrate specificity and structural relationships of the natural substrates trichohyalin and filaggrin. J. Biol. Chem. 271, 30709–16 (1996).

13. Musse, A. A. et al. Peptidylarginine deiminase 2 (PAD2) overexpression in transgenic mice leads to myelin loss in the central nervous system. DMM Dis. Model. Mech. 1, 229–240 (2008).

14. Christophorou, M. A. et al. Citrullination regulates pluripotency and histone H1 binding to chromatin. Nature 507, 104–108 (2014).

15. Deplus, R. et al. Citrullination of DNMT3A by PADI4 regulates its stability and controls DNA methylation. Nucleic Acids Res. 42, 8285–8296 (2014).

16. Falcão, A. M. et al. PAD2-Mediated Citrullination Contributes to Efficient Oligodendrocyte Differentiation and Myelination. Cell Rep. 27, 1090–1102.e10 (2019).

17. Méchin, M. C., Takahara, H. & Simon, M. Deimination and peptidylarginine deiminases in skin physiology and diseases. Int. J. Mol. Sci. 21, 1–15 (2020).

18. Zhang, X. et al. Peptidylarginine deiminase 1-catalyzed histone citrullination is essential for early embryo development OPEN. Sci. Rep. 6, (2016).

19. Kholia, S., et al. A novel role for peptidylarginine deiminases in microvesicle release reveals therapeutic potential of PAD inhibition in sensitizing prostate cancer cells to chemotherapy. J. Extracell. Vesicles 4, (2015).

20. Haider, L. et al. The topograpy of demyelination and neurodegeneration in the multiple sclerosis brain. Brain 139, 807–815 (2016).

21. Sokolove, J. et al. Citrullination within the atherosclerotic plaque: A potential target for the anti-citrullinated protein antibody response in rheumatoid arthritis. Arthritis Rheum. 65, 1719–1724 (2013).

22. Khandpur, R. et al. NETs Are a Source of Citrullinated Autoantigens and Stimulate Inflammatory Responses in Rheumatoid Arthritis. Sci. Transl. Med. 5, (2013).

23. Corsiero, E., Pratesi, F., Prediletto, E., Bombardieri, M. & Migliorini, P. NETosis as source of autoantigens in rheumatoid arthritis. Front. Immunol. 7, 1–9 (2016).

24. Singh, U., Singh, S., Singh, N. K., Verma, P. K. & Singh, S. Anticyclic citrullinated peptide autoantibodies in systemic lupus erythematosus. Rheumatol. Int. 31, 765–767 (2011).

25. Gupta, S. & Kaplan, M. J. The role of neutrophils and NETosis in autoimmune and renal diseases. Nat. Rev. Nephrol. 12, 402–413 (2016).

26. Schellekens, G. A., de Jong, B. A., van den Hoogen, F. H., van de Putte, L. B. & van Venrooij, W. J. Citrulline is an Essential Constituent of Antigenic Determinants Recognized by Rheumatoid Arthritis − specific Autoantibodies. J Immunol 195, 8–16 (1998).

27. Suzuki, A. et al. Functional haplotypes of PADI4, encoding citrullinating enzyme peptidylarginine deiminase 4, are associated with rheumatoid arthritis. Nat. Genet. 34, 395– 402 (2003).

28. Yamada, R., Suzuki, A., Chang, X. & Yamamoto, K. Peptidylarginine deiminase type 4: Identification of a rheumatoid arthritis-susceptible gene. Trends Mol. Med. 9, 503–508 (2003).

29. Lee, Y. H. & Bae, S. C. Association between susceptibility to rheumatoid arthritis and PADI4 polymorphisms: A meta-analysis. Clin. Rheumatol. 35, 961–971 (2016).

30. Christophorou, M. A. The virtues and vices of protein citrullination. R. Soc. open Sci. 9, (2022).

31. Ciesielski, O. et al. Citrullination in the pathology of inflammatory and autoimmune disorders: recent advances and future perspectives. 79, 94 (2022).

32. Fert-Bober, J. et al. Mapping Citrullinated Sites in Multiple Organs of Mice Using Hypercitrullinated Library. J. Proteome Res. (2019). doi:10.1021/acs.jproteome.9b00118

33. Bekker-Jensen, D. B. et al. An Optimized Shotgun Strategy for the Rapid Generation of Comprehensive Human Proteomes. Cell Syst. (2017). doi:10.1016/j.cels.2017.05.009

34. Nakashima, K. et al. Molecular Characterization of Peptidylarginine Deiminase in HL60 Cells Induced by Retinoic Acid and 1a,25-Dihydroxyvitamin D 3. J. Biol. Chem. 274, 27786– 27792 (1999).

35. Mondal, S. & Thompson, P. R. Protein Arginine Deiminases (PADs): Biochemistry and Chemical Biology of Protein Citrullination. Acc. Chem. Res. 52, 818–832 (2019).

36. Lee, C.-Y. et al. Mining the Human Tissue Proteome for Protein Citrullination*. Mol. Cell. Proteomics 17, 1378–1391 (2018).

37. Kelstrup, C. D., Young, C., Lavallee, R., Nielsen, M. L. & Olsen, J. V. Optimized fast and sensitive acquisition methods for shotgun proteomics on a quadrupole orbitrap mass spectrometer. J. Proteome Res. 11, 3487–3497 (2012).

38. Sylvestersen, K. B., Young, C. & Nielsen, M. L. Advances in characterizing ubiquitylation sites by mass spectrometry. Curr. Opin. Chem. Biol. 17, 49–58 (2013).

39. Cox, J. et al. Accurate proteome-wide label-free quantification by delayed normalization and maximal peptide ratio extraction, termed MaxLFQ. Mol. Cell. Proteomics 13, 2513–26 (2014).

40. Alyami, H. M. et al. Role of NOD1/NOD2 receptors in Fusobacterium nucleatum mediated NETosis. Microb. Pathog. 131, 53–64 (2019).

41. Wang, Y. et al. Histone hypercitrullination mediates chromatin decondensation and neutrophil extracellular trap formation. J. Cell Biol 184, 205–213 (2009).

42. Kelstrup, C. D. et al. Rapid and deep proteomes by faster sequencing on a benchtop quadrupole ultra-high-field orbitrap mass spectrometer. J. Proteome Res. 13, 6187–6195 (2014).

43. Foulquier, C. et al. Peptidyl arginine deiminase type 2 (PAD-2) and PAD-4 but not PAD-1, PAD-3, and PAD-6 are expressed in rheumatoid arthritis synovium in close association with tissue inflammation. Arthritis Rheum. 56, 3541–3553 (2007).

44. Zhou, Y. et al. Spontaneous Secretion of the Citrullination Enzyme PAD2 and Cell Surface Exposure of PAD4 by Neutrophils. Front. Immunol. 8, 1200 (2017).

45. Lakschevitz, F. S. et al. Identification of neutrophil surface marker changes in health and inflammation using high-throughput screening flow cytometry. Exp. Cell Res. 342, 200–209 (2016).

46. Eldewi, D. M. et al. Expression levels of complement regulatory proteins (CD35, CD55 and CD59) on peripheral blood cells of patients with chronic kidney disease. Int. J. Gen. Med. 12, 343–351 (2019).

47. Chick, J. M. et al. An ultra-tolerant database search reveals that a myriad of modified peptides contributes to unassigned spectra in shotgun proteomics HHS Public Access Author manuscript. Nat Biotechnol 33, 743–749 (2015).

48. Hao, G. et al. Neutral Loss of Isocyanic Acid in Peptide CID Spectra: A Novel Diagnostic Marker for Mass Spectrometric Identification of Protein Citrullination. JAM 20, 723–727 (2008).

49. Olsen, J. V et al. Quantitative Phosphoproteomics Reveals Widespread Full Phosphorylation Site Occupancy During Mitosis. Sci. Signal. 3, (2010).

50. Hansen, B. K. et al. Analysis of human acetylation stoichiometry defines mechanistic constraints on protein regulation. Nat. Commun. 10, (2019).

51. Zielinska, D. F., Gnad, F., Wiśniewski, J. R. & Mann, M. Precision mapping of an in vivo N-glycoproteome reveals rigid topological and sequence constraints. Cell 141, 897–907 (2010).

52. Guo, Q., Bedford, M. T. & Fast, W. Discovery of Peptidylarginine Deiminase-4 Substrates by Protein Array: Antagonistic Citrullination and Methylation of Human Ribosomal Protein S2. Mol. Biosyst. 7, 2286–95 (2011).

53. Nakashima, K., Hagiwara, T. & Yamada, M. Nuclear localization of peptidylarginine deiminase V and histone deimination in granulocytes. J. Biol. Chem. 277, 49562–49568 (2002).

54. Bekker-Jensen, D. B. et al. A compact quadrupole-orbitrap mass spectrometer with FAIMS interface improves proteome coverage in short LC gradients. Mol. Cell. Proteomics 19, 716– 729 (2020).

55. Wang, D. et al. AAgAtlas 1.0: A human autoantigen database. Nucleic Acids Res. 45, D769–D776 (2017).

56. Lewis, H. D. et al. Inhibition of PAD4 activity is sufficient to disrupt mouse and human NET formation. (2015). doi:10.1038/nCHeMBIO.1735

57. Mondal, S. et al. Site-specific incorporation of citrulline into proteins in mammalian cells. Nat. Commun. 12, (2021).

58. Kinloch, A. et al. Identification of citrullinated alpha-enolase as a candidate autoantigen in rheumatoid arthritis. Arthritis Res. Ther. 7, 1421–1429 (2005).

59. Olsson, T. et al. Increased numbers of T cells recognizing multiple myelin basic protein epitopes in multiple sclerosis. Eur. J. Immunol. 22, 1083–1087 (1992).

60. Hagiwara, T., Nakashima, K., Hirano, H., Senshu, T. & Yamada, M. Deimination of Arginine Residues in Nucleophosmin/B23 and Histones in HL60 Granulocytes. Biochem. Biophys. Res. Commun. 290, 979–983 (2002).

61. Sohn, D. H. et al. Local joint inflammation and histone citrullination in a murine model of the transition from preclinical autoimmunity to inflammatory arthritis. Arthritis Rheumatol. 67, 2877–2887 (2015).

62. Cherrington, B. D. et al. Potential Role for PAD2 in Gene Regulation in Breast Cancer Cells. PLoS One 7, e41242 (2012).

63. Cuthbert, G. L. et al. Histone deimination antagonizes arginine methylation. Cell 118, 545– 553 (2004).

64. Leshner, M. et al. PAD4 mediated histone hypercitrullination induces heterochromatin decondensation and chromatin unfolding to form neutrophil extracellular trap-like structures. Front. Immunol. 3, 1–11 (2012).

65. Lee, Y.-H., Coonrod, S. A., Kraus, W. L., Jelinek, M. A. & Stallcup, M. R. Regulation of coactivator complex assembly and function by protein arginine methylation and demethylimination. Proc. Natl. Acad. Sci. U. S. A. 102, 3611–6 (2005).

66. Wang, S. & Wang, Y. Peptidylarginine deiminases in citrullination, gene regulation, health and pathogenesis. Biochim. Biophys. Acta 1829, 1126–35 (2013).

67. Sharma, P. et al. Arginine Citrullination at the C-Terminal Domain Controls RNA Polymerase II Transcription. Mol. Cell 73, 84–96.e7 (2019).

68. Wang, Y. et al. Human PAD4 regulates histone arginine methylation levels via demethylimination. Science (80-.). 306, 279–283 (2004).

69. Zhai, Q., Wang, L., Zhao, P. & Li, T. Role of citrullination modification catalyzed by peptidylarginine deiminase 4 in gene transcriptional regulation. Acta Biochim. Biophys. Sin. (Shanghai). 49, 567–572 (2017).

70. Fuhrmann, J. & Thompson, P. R. Protein Arginine Methylation and Citrullination in Epigenetic Regulation. ACS Chem. Biol. 11, 654–68 (2016).

71. Clancy, K. W. et al. Citrullination/Methylation Crosstalk on Histone H3 Regulates ER-Target Gene Transcription. ACS Chem. Biol. 12, 1691–1702 (2017).

72. Tanikawa, C. et al. Regulation of protein citrullination through p53/PADI4Network in DNA damage response. Cancer Res. 69, 8761–8769 (2009).

73. Schreiber, V., Molinete, M., Boeuf, H., de Murcia, G. & Ménissier-de Murcia, J. The human poly(ADP-ribose) polymerase nuclear localization signal is a bipartite element functionally separate from DNA binding and catalytic activity. EMBO J. 11, 3263–3269 (1992).

74. Doll, S. G. et al. Recognition of the TDP-43 nuclear localization signal by importin α1/β. Cell Rep. 39, (2022).

75. Psakhye, I. & Jentsch, S. Protein group modification and synergy in the SUMO pathway as exemplified in DNA repair. Cell 151, 807–820 (2012).

76. Huang, D. & Kraus, W. L. The expanding universe of PARP1-mediated molecular and therapeutic mechanisms. Mol. Cell 82, 2315–2334 (2022).

77. Lewis, H. D. & Nacht, M. IPAD or PADi - ‘tablets’ with therapeutic disease potential? Curr. Opin. Chem. Biol. 33, 169–178 (2016).

78. Lange, S. et al. Peptidylarginine Deiminases-Roles in Cancer and Neurodegeneration and Possible Avenues for Therapeutic Intervention via Modulation of Exosome and Microvesicle (EMV) Release? Int. J. Mol. Sci. 18, (2017).

79. Gudmann, N. S., Hansen, N. U. B., Jensen, A. C. B., Karsdal, M. A. & Siebuhr, A. S. Biological relevance of citrullinations: diagnostic, prognostic and therapeutic options. Autoimmunity 48, 73–79 (2015).

80. Matsuo, K. et al. Identification of novel citrullinated autoantigens of synovium in rheumatoid arthritis using a proteomic approach. Arthritis Res. Ther. 8, R175 (2006).

81. Li, P. et al. PAD4 is essential for antibacterial innate immunity mediated by neutrophil extracellular traps. J. Exp. Med. 207, 1853–62 (2010).

82. Harding, C. R. & Scott, I. R. Histidine-rich proteins (filaggrins): Structural and functional heterogeneity during epidermal differentiation. J. Mol. Biol. 170, 651–673 (1983).

83. Méchin, M. C. et al. Update on peptidylarginine deiminases and deimination in skin physiology and severe human diseases. Int. J. Cosmet. Sci. 29, 147–168 (2007).

84. Bernstein, B. E. et al. Genomic maps and comparative analysis of histone modifications in human and mouse. Cell 120, 169–181 (2005).

85. Zhang, X., Gamble, M. J., Stadler, S. ¤, Cherrington, B. D. & Causey, C. P. Genome-Wide Analysis Reveals PADI4 Cooperates with Elk-1 to Activate c-Fos Expression in Breast Cancer Cells. PLoS Genet 7, 1002112 (2011).

86. Schölz, C. et al. Acetylation site specificities of lysine deacetylase inhibitors in human cells. Nat. Biotechnol. 33, 415–425 (2015).

87. Perez-Riverol, Y. et al. The PRIDE database resources in 2022: A hub for mass spectrometry-based proteomics evidences. Nucleic Acids Res. 50, D543–D552 (2022).

88. Senshu, T., Sato, T., Inoue, T., Akiyama, K. & Asaga, H. Detection of citrulline residues in deiminated proteins on polyvinylidene difluoride membrane. Anal. Biochem. 203, 94–100 (1992).

89. Cox, J. & Mann, M. Quantitative, High-Resolution Proteomics for Data-Driven Systems Biology. (2011). doi:10.1146/annurev-biochem-061308-093216

90. Cox, J. & Mann, M. MaxQuant enables high peptide identification rates, individualized p.p.b.-range mass accuracies and proteome-wide protein quantification. Nat. Biotechnol. 26, 1367–1372 (2008).

91. Tyanova, S. et al. The Perseus computational platform for comprehensive analysis of (prote)omics data. Nature Methods (2016). doi:10.1038/nmeth.3901

92. Sherman, B. T., et al. DAVID: a web server for functional enrichment analysis and functional annotation of gene lists (2021 update). Nucleic Acids Res. 50, W216–W221 (2022).

93. Huang, D. W., Sherman, B. T. & Lempicki, R. A. Systematic and integrative analysis of large gene lists using DAVID bioinformatics resources. Nat. Protoc. 4, 44–57 (2009).

94. Hulsen, T., de Vlieg, J. & Alkema, W. BioVenn - A web application for the comparison and visualization of biological lists using area-proportional Venn diagrams. BMC Genomics 9, 1– 6 (2008).

95. Spitzer, M., Wildenhain, J., Rappsilber, J. & Tyers, M. BoxPlotR: a web tool for generation of box plots. Nat. Methods 2014 112 11, 121–122 (2014).

96. Horn, H. et al. KinomeXplorer: An integrated platform for kinome biology studies. Nature Methods 11, 603–604 (2014).

97. Linding, R. et al. NetworKIN: A resource for exploring cellular phosphorylation networks. Nucleic Acids Res. 36, D695 (2008).

98. Ham, A.-J L, et al. Improved visualization of protein consensus sequences by iceLogo. Nat. Methods 2009 611 6, 786–787 (2009).

99. Puente-Santamaria, L., Wasserman, W. W. & Del Peso, L. TFEA.ChIP: A tool kit for transcription factor binding site enrichment analysis capitalizing on ChIP-seq datasets. Bioinformatics 35, 5339–5340 (2019).

100. Szklarczyk, D., et al. STRING v11: Protein-protein association networks with increased coverage, supporting functional discovery in genome-wide experimental datasets. Nucleic Acids Res. 47, D607–D613 (2019).

101. Mészáros, B., Erdös, G. & Dosztányi, Z. IUPred2A: Context-dependent prediction of protein disorder as a function of redox state and protein binding. Nucleic Acids Res. 46, W329– W337 (2018).

102. Bateman, A. et al. UniProt: The universal protein knowledgebase in 2021. Nucleic Acids Res. 49, D480–D489 (2021).

103. Meyer, J. G. et al. Quantification of Lysine Acetylation and Succinylation Stoichiometry in Proteins Using Mass Spectrometric Data-Independent Acquisitions (SWATH). J. Am. Soc. Mass Spectrom. 27, 1758–1771 (2016).

104. Wu, R. et al. A large-scale method to measure absolute protein phosphorylation stoichiometries. Nat. Methods 8, 677–683 (2011).

105. Sun, S. & Zhang, H. Large-Scale Measurement of Absolute Protein Glycosylation Stoichiometry. Anal. Chem. 87, 6479–6482 (2015).

